# Template switching by coronavirus polymerase requires helicase activity and is stimulated by remdesivir and molnupiravir

**DOI:** 10.1101/2025.10.03.680419

**Authors:** Asif Rakib, Arnab Das, Subhas C. Bera, Pim P. B. America, Misha Klein, Thomas K. Anderson, John C. Marecki, Bing Wang, Eline Bogers, Joy Y. Feng, John P. Bilello, Flávia S. Papini, Quinte Smitskamp, Jamie J. Arnold, Irina Artsimovitch, Robert N. Kirchdoerfer, Craig E. Cameron, Kevin D. Raney, David Dulin

## Abstract

Polymerase template switching is an essential mechanism in coronaviruses (CoVs) that enables both sub-genomic (sg) RNA synthesis and increases genomic diversity via RNA recombination. Despite its importance, the molecular mechanism of CoV polymerase template switching remains unclear. Using magnetic tweezers, we show that the CoV non-structural protein (nsp) 13-helicase drives polymerase template switching, followed by copy-back RNA synthesis. This activity requires nsp13-helicase ATPase activity and a duplex RNA downstream of the CoV polymerase. This novel function of nsp13-helicase is targeted by the nucleotide analogs remdesivir and molnupiravir, whose incorporation in the nascent strand increases CoV polymerase template switching probability, leading to defective RNA production. We propose a novel mechanism of action where incorporation of these analogs dramatically reduces full length genome copy number by stimulating polymerase template switching. Our study further demonstrates nsp13-helicase’s central role in CoV replication and how this enzyme function can be indirectly targeted by analogs.

## Introduction

The SARS-CoV-2 pandemic has shown how vulnerable we are to viral pandemics without available therapeutics. Although effective drugs against CoVs are now available, developing new broad-spectrum antivirals remains a research priority in preparation for future SARS-CoV-2 variants and other CoV zoonose.

CoVs possess a ~30 kb, positive-sense, single-stranded (ss) RNA genome. The 5’-proximal two-thirds encode for sixteen non-structural proteins (nsps), while the 3’-proximal one-third encodes for structural proteins and accessory proteins (*1*). Nsps are directly translated from the RNA genome and the structural proteins are expressed from a set of 3’ co-terminal nested sub-genomic messenger RNAs that are also 5’ co-terminal nested (*2*). The sub-genomic negative-strand RNAs (sgRNAs) are produced during negative-strand synthesis (*3–5*). It has been proposed to occur via a mechanism similar to homology assisted copy-choice RNA recombination and requires the CoV polymerase to jump ~20 kb from transcription regulatory sequence (TRS) ending each structural protein gene to the TRS-leader at the 5’-UTR and switch template (*2*). These sgRNAs are subsequently used as a template for synthesis of subgenomic messenger (sgm) RNAs. The CoV polymerase is part of the replication-transcription complex (RTC) that synthesizes all viral RNAs in the infected cell. The CoV RTC is made of a core, i.e. nsp7, nsp8 and nsp12-polymerase in a 1:2:1 stoichiometry, which associates with several other nsps, including nsp13-helicase (*6–15*). Nsp13-helicase is a superfamily 1B helicase that unwinds double-stranded (ds) nucleic acids with a 5’ to 3’ polarity (*16–18*). Structural studies revealed that two nsp13-helicases binds with the core RTC, i.e. nsp13.1 and nsp13.2, and nsp13.1 has been shown to associate with the template RNA (*19–22*). The CoV nsp13-helicase has been proposed to carry out several functions within the RTC, such as assisting the RNA synthesis through secondary RNA structures and facilitating RTC backtracking, which is suggested to be required for proofreading and polymerase strand switching (*23*). Using high-throughput magnetic tweezers, we recently demonstrated that nsp13.2 assists RNA synthesis through duplex RNA by translocating on the strand opposite to the template (*24*). Nsp13.1 has been shown to associate with the template RNA and has therefore been proposed to promote RTC backtracking, though this function has not been directly demonstrated (*20*).

The RTC is structurally and functionally conserved amongst coronaviruses, making it an important target for broad-spectrum antivirals (*25*), such as nucleoside analogs, whose mechanisms of action have been shown to induce pausing, termination, or mutations during viral RNA synthesis. Remdesivir (RDV) is an adenosine analog with a 1’-cyano modification and was the first antiviral approved to treat SARS-CoV-2 infection (*26–30*). Upon incorporation into the nascent RNA, RDV-triphosphate (RDV-TP) induces long pauses that have been linked to polymerase backtracking due to a steric clash between RDV 1’-cyano moiety and serine-861 of nsp12-polymerase (*10, 31–33*). The 5’-monophosphate (MP) of RDV (RDV-MP) insertion in the template strand has been shown to halt RTC elongation (*34*). Molnupiravir (MPV), a CTP analog with Emergency Use Authorization (EUA) to treat SARS-CoV-2 infection, has shown efficacy against SARS-CoV-2 (*35–37*) and several other RNA viruses (*38–44*). MPV has been reported to induce mutations via the template strand following initial incorporation during negative-strand synthesis (*45, 46*). The currently accepted mechanisms of actions of MPV and RDV have been characterized using the core RTC only, i.e. without other nsp’s.(*45, 47*). Because nsp13-helicase plays a major role in the RTC elongation dynamics through duplex RNA (*24*), we hypothesize that it may affect the mechanism of action of antivirals.

Here, we employ high-throughput magnetic tweezers to investigate the function of nsp13-helicase in complex with an elongating RTC, and to determine whether the presence of nsp13-helicase affects the mechanism of action of several nucleotide analogs targeting the CoV polymerase. We show that the CoV RTC switches template intramolecularly to perform copy-back RNA synthesis and that nsp13-helicase ATPase activity and a dsRNA fork downstream of the polymerase are both required for this activity to occur. These requirements suggest a polymerase backtracking-dependent mechanism, further potentiated by nsp13.1 ATPase activity. Decreasing NTP concentration increases copy-back RNA synthesis probability, and this effect is more potent with ATP. Investigating the mechanism of action of RDV, we show that adding nsp13-helicase entirely removes the RDV-TP-induced pauses, demonstrating that nsp13-helicase enables the elongating RTC to overcome the RDV-TP incorporation barrier to elongation. We show that RDV-TP incorporation increases the probability of intramolecular template switching and copy-back RNA synthesis, reducing CoV RTC processivity even at low concentration. A similar mode of action was also observed with MPV. In contrast, when using arabinose-UTP – an analog that induces a long-lived pause following incorporation –, we observed a decrease in the probability of copy-back RNA synthesis and a further stabilization of the long-lived pause leading a decrease in RTC processivity.

We propose a model where MPV-TP and RDV-TP incorporation increases the probability for nsp13.2 to disengage from the non-template RNA, while nsp13.1 engages with the template RNA and pushes the RTC into a deep backtrack. The 3’-end of the product RNA can thereafter snap back and self-anneal, followed by intramolecular polymerase template switching and copy-back RNA synthesis. We propose that such activity in vivo could significantly decrease the fraction of full-length viral genome, offering another explanation for the potency of these antivirals. Analogs targeting the viral polymerase can have unpredictable effects on the function of associated cofactors, further highlighting the importance of investigating their mechanisms of action in the context of a complete replication complex.

## Results

### Nsp13-helicase induces SARS-CoV-2 RTC template-strand switching

To investigate the elongation dynamics of SARS-CoV-2 core RTC associated with nsp13-helicase, we employed a high-throughput magnetic tweezers assay (**Figure 1A**) (*48–50*). Briefly, magnetic beads were tethered to the glass surface of a flow cell with an RNA construct (**Materials and Methods, Figure S1**). When using a dsRNA construct (**Figure S1**), the SARS-CoV-2 RTC assembled at the short hairpin terminating the template strand at the 3’-end (**Figure 1A**). The RTC elongated the stem of the short hairpin, which concomitantly converted the dsRNA tether into ssRNA and increased the tether extension (**Figure 1A**) (*51*). Similarly, when using a ssRNA template, the tether extension decreased during the conversion of the ssRNA template into dsRNA by the CoV RTC (**Figure S2A**). Finally, when using a closed hairpin construct, the tether extension increased when the RTC elongated and opened the hairpin (**Figure S2E**). For these three constructs, we followed the tridimensional position of the magnetic bead to measure the change in extension of the tether and extract the length of the product RNA strand (**Materials and Methods**)

**Figure 1:**
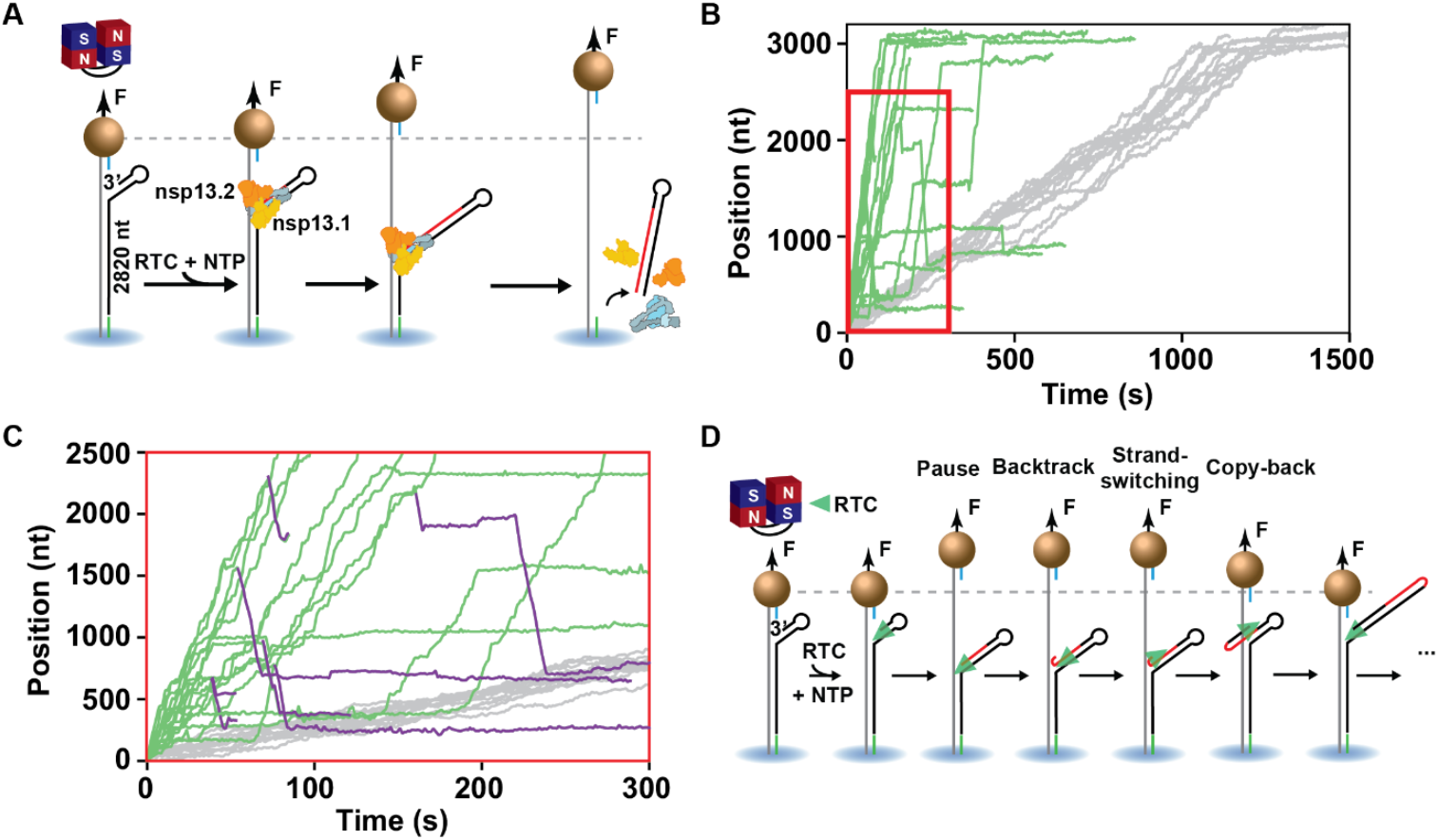
The SARS-CoV-2 nsp13-helicase induces RTC reversal events. **(A)** Schematic of the magnetic tweezers assay to monitor RTC elongation on dsRNA. The core RTC (nsp7, nsp8 & nsp12) and the nsp13-helicase assembles to the small hairpin terminating the 3’ extremity of the template, followed by elongation in the presence of NTP. **(B)** RNA synthesis activity traces obtained with (green) (*N* = 17) or without (gray) (*N* = 13) 20 nM nsp13-helicase. The NTPs were maintained at concentration of 500 µM. **(C)** A zoomed-in view of the elongation traces highlighted by the red rectangle in **(B)**, showing RNA synthesis in the reversed direction (violet). **(D)** Model describing the mechanism behind CoV RTC reversals.

We recently showed that, in the presence of nsp13-helicase, a CoV RTC elongating through a duplex RNA is on average ~10-times faster than the core RTC alone (**Figure 1B**) (*24*). Surprisingly, a significant fraction of SARS-CoV-2 RTC reversed direction in a processive manner, i.e. over tens to hundreds of nucleotides distance (**Figure 1C**), precluding a diffusive process such as polymerase backtracking (*52, 53*). We coin the term “reversals” to describe these events. Reversals were not observed when using either the core RTC alone (**Figure 1B**) or when using a ssRNA template (**Figure S2AB**). A possible origin for the reversals could be a nick in the template strand (**Figure 2SCD**). In such a case, the RTC would have elongated along the template strand until it encounters the nick. The RTC would have then fallen off, followed by another RTC assembling at the 3’-end of the remaining template strand and elongating it. The conversion of the ssRNA part of the tether into dsRNA would have reduced the tether extension (**Figure 2SD**), similarly to what occured when using the ssRNA template (**Figure S2A**). To verify this hypothesis, we used a closed hairpin instead of a dsRNA construct. If the hairpin stem was nicked, the bead would have been lost when the RTC elongated through the nick, and we should have therefore not been able to monitor reversals. However, reversals were captured when using a hairpin construct (**Figure S2EF**), arguing that these events did not result from a nick in the template strand. The probability of reversals decreased by 10-fold when increasing the force applied to the hairpin from 9 pN to 20 pN (**Figure S2G, Table S1**). We previously showed that the core RTC had a higher probability to enter a backtrack state when lowering the force applied to the RNA hairpin, which increased the stability of the dsRNA fork (*52*). We propose that backtracking is a necessary intermediate for a reversal to occur, consistent with what has been previously shown for other viral RNA-dependent RNA polymerases (RdRps) (*54, 55*). Furthermore, enterovirus A-71 and poliovirus RdRps reversals were shown to originate from polymerase backtracking followed by intramolecular polymerase template switching and copy-back RNA synthesis (*55*). CoV RTC template switching further required nsp13-helicase ATPase activity, as employing ATPase-dead mutant K288A nsp13-helicase instead resulted in very slow traces and no reversals (**Figure S2H**).

**Figure 2:**
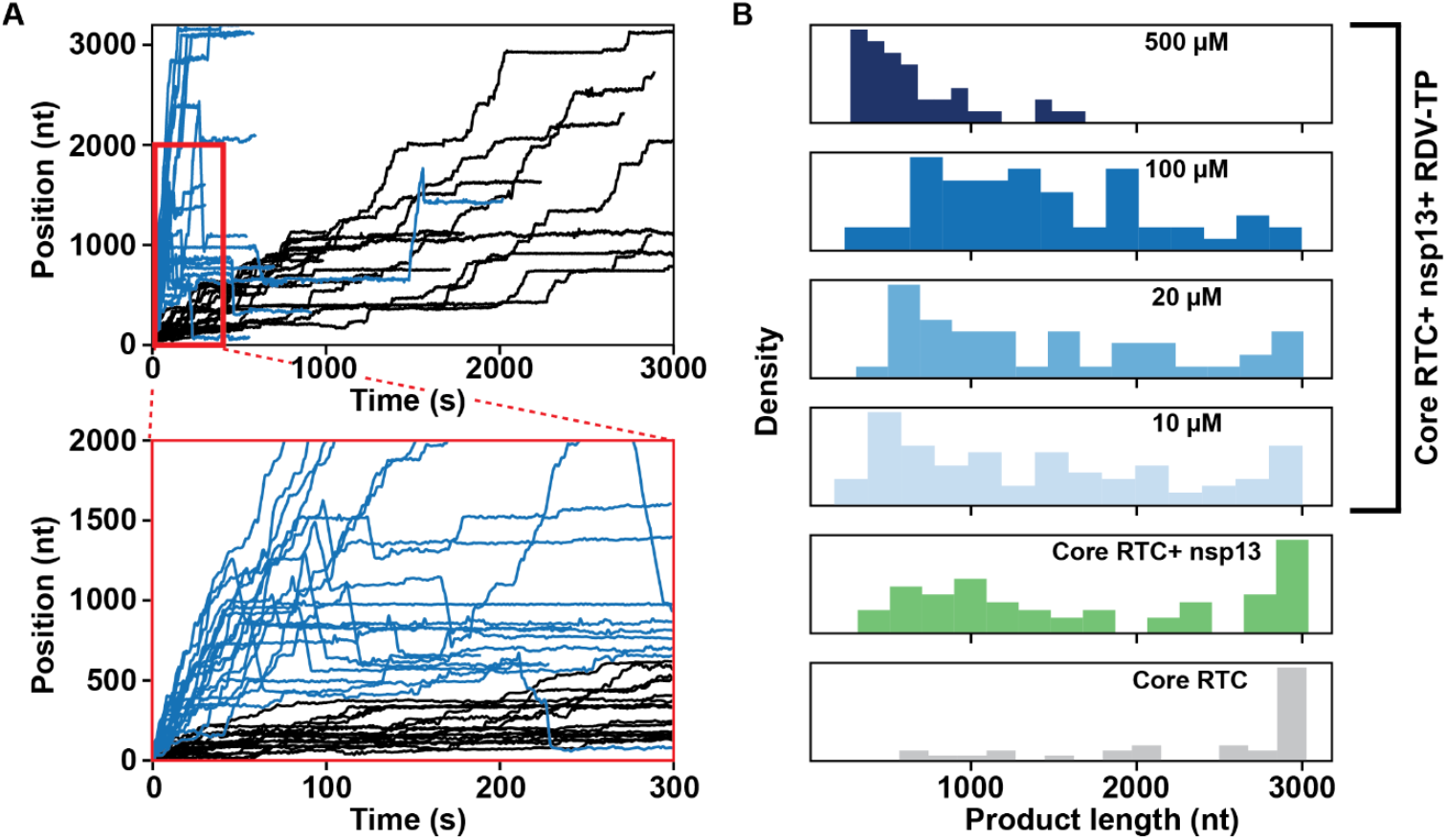
Remdesivir incorporation increases reversals probability. **(A)** Elongation traces obtained with (blue) or without (black) 20 nM nsp13-helicase, 100 µM RDV-TP and 500 µM NTPs. The bottom panel represents a zoomed-in view of the elongation traces highlighted with the red rectangle. **(B)** Histogram of the forward processivity before the first reversal at different experimental conditions: without nsp13-helicase (gray), with 20 nM nsp13-helicase (green), with 20 nM nsp13-helicase and the indicated concentration of RDV-TP (10 µM, 20 µM, 100 µM, 500 µM) shown in a gradient from light blue to dark blue. All the experiments were performed in the presence of 500 µM NTP, at 20 pN and 25 °C.

We previously reported that low NTP concentrations increases the probability for SARS-CoV-2 core RTC to enter the slow nucleotide addition pathways (*52*), which can branch out into the long-lived pauses and backtrack states (*24*). Therefore, lowering any of the NTP concentrations may stimulate reversals. When decreasing a single NTP to 50 µM while keeping the others at 500 µM, we saw an increase in reversals probability compared to when all NTP are at 500 µM (**Figure S3**). The RTC was particularly sensitive to ATP concentration. Indeed, at low ATP, reversals were detected very early in the trace with respect to the template strand length (~2.9 kb) and none of the elongating RTC reaches the end of the template (**Figure S3**). We next investigated whether nsp13-helicase affinity for a specific NTP could explain such dependence of the reversals to specific NTP (**Figure S4A, Figure S1C**). Nsp13-helicase exhibits an unwinding velocity of (328 ± 5) bp/s at 1 mM ATP, which decreases to (198 ± 5) bp/s and (147 ± 5) bp/s at 1 mM UTP and CTP, respectively (**Figure S4EGI**). The ATP-dependent value is in good agreement with previous measurements done with optical tweezers (*16*). Fitting a Michaelis-Menten model (**Materials and Methods**) to the unwinding velocity for each NTP, we determined a *K*_*m*_ ranging from 37 µM (ATP) to 88 µM (GTP) (**Figure S5**). The *V*_*max*_ was notably higher for ATP and GTP (~400 bp/s) compared to CTP and UTP (~200 bp/s), indicating a faster cycle for purine over pyrimidine. Additionally, the lower *K*_*m*_ for ATP, as compared to GTP, suggests a higher affinity for ATP. These results are largely in agreement with a previous bulk measurement on human coronavirus 229E nsp13-helicase (*56*). The high NTP hydrolysis rate, even without ssRNA (*57*), and the high affinity of nsp13-helicase with ATP suggest that ATP may be depleted faster than the other NTP by the helicase to a very low concentration, further increasing the probability for the RTC to enter a reversal.

Previous studies showed that nsp13.1 and nsp13.2 allosterically control each other’s productive engagement with either the template or the non-template strand, respectively, allowing only one helicase to actively translocate at a time (*21, 24*). We propose a model where the RTC switches from a nsp13.2-to nsp13.1-engaged state (*24*). Following entry into backtrack state, the RTC is further pushed backward by nsp13.1, enabling the product RNA 3’-end to snap back and self-anneal. The RTC then switches template to perform copy-back RNA synthesis, using the former product strand as a template (**Figure 1D**).

### Remdesivir incorporation increases reversal occurrence

We then investigated how having a single RDV inserted in the middle of a ~1 kb long single-stranded template RNA could inhibit an elongating core RTC (**Materials and Methods, Figure S1D**). In the presence of 500 µM NTP, we observed no significant pause at the ~500 nt position (**Figure S6B**). A previous study has reported that decreasing UTP concentration increases the pausing effect of a single RDV-MP inserted in the template strand (*34*). Therefore, we repeated the experiment at 50 µM UTP, keeping all other NTPs at 500 µM, but no significant pause was observed at the RDV-MP position in the template strand (**Figure S6C**). Our assay is able to detect a pause induced by a single modified nucleotide in the template strand, as we show here for 2’-O-Me-AMP (**Figure S1D** and **Figure S6D**), where most traces are blocked for a duration exceeding the experiments (30 min) at the expected position on the template strand (**Figure S6D**). This result is consistent with a previous study reporting termination by 2’-O-Me-AMP insertion (*58*). We conclude that RDV’s mechanism of action is likely not to pause or block an elongating RTC when present in the template strand.

We and others showed that RDV-TP incorporation in the nascent strand induces backtracking-related pauses to an elongating core RTC (*10, 31–33*). Here, we sought to determine whether such pauses remain in the presence of nsp13-helicase. We first investigated the impact of RDV on a core RTC elongating on a dsRNA template. With 100 µM RDV-TP and 500 µM NTPs, we observed a massive increase in the pause duration and frequency (**Figure 2A** and **Figure S7A**). However, in the presence of nsp13-helicase, the long-lived pauses almost completely disappeared from the traces (**Figure 2A**). To further analyze the trace, we performed a dwell time analysis, where we scan the traces with a non-overlapping 10 nt window to measure the duration of ten successive nucleotide addition cycles (*24, 59*). A dwell time is, therefore, a measure of the biochemical event that rate-limits the ten successive nucleotide addition cycles. The resulting dwell times were assembled into a probability density distribution represented in a log-binned histogram (**Figure S7B**) (*59*). The core RTC elongation dynamic can be decomposed into four different distributions, i.e. a gamma distribution for the fast nucleotide addition (FNA) pathway, two exponential distributions for the slow and very slow nucleotide addition (SNA and VSNA, respectively) pathways, and a power law with *t*^−3/2^ exponent for the long-lived pauses (LLP) (**Materials and Methods, Figure S7C**). In the presence of nsp13-helicase, another gamma distribution appears on the left of the FNA gamma distribution and describes the very fast nucleotide addition (VFNA, **Figure S8AB**) pathway. Here, nsp13.2 employs NTP hydrolysis to assist the RTC elongating through the duplex RNA fork (*24*). All the distributions are fitted at once using a maximum likelihood estimation procedure to extract the probability and characteristic timescale of each distribution (**Materials and Methods**). In the absence of nsp13-helicase, only the LLP probability increases significantly with RDV-TP concentration. Indeed, RDV-TP incorporation induces LLP related to polymerase backtracking upon incorporation by the core RTC (**Figure S7DE, Table S2**). The FNA characteristic timescale is unaffected by RDV-TP concentration, further indicating that RDV-TP is likely not incorporated in this pathway (*49*). The presence of the dsRNA fork further slows down the core RTC elongation in comparison to the ssRNA template (*49*). In the presence of nsp13-helicase, the distributions are largely unchanged for RDV-TP concentration up to 100 µM (**Figure S8A**), indicating that nsp13-helicase largely removes the barrier to translocation induced by a single RDV-TP incorporation (*32*). However, at 500 µM RDV-TP, the VFNA gamma distribution almost entirely disappears in favor of the FNA gamma distribution (**Figure S8A**), effectively nullifying nsp13.2’s ability to assist the RTC to elongate through the duplex RNA. RDV-TP is a poor substrate for nsp13-helicase and is therefore unlikely to be used as a substitute for ATP (**Figure S4F**). However, RDV-TP is an excellent substrate for nsp12-polymerase (*49, 60*) and could be inserted through multiple successive nucleotide addition cycles when present at a high concentration. The barrier induced by multiple successive RDV-TP incorporations is much higher than for a single incorporation (*31*), and nsp12-polymerase is unlikely to overcome it even when assisted by nsp13-helicase.

While we do not notice a significant change in the nucleotide addition dynamics of the SARS-CoV-2 RTC until reaching a RDV-TP concentration above 100 µM, i.e. two order of magnitude higher than the measured intracellular effective concentration of RDV-TP (2-4 µM) (*61*), the probability of reversals increases dramatically even at low concentration (**Figure 2A**). The number of reversals increases by almost 50%, i.e. from (0.44 ± 0.13) bp/s to (0.64 ± 0.1) bp/s when increasing RDV-TP concentration from 0 to 10 µM (**Figure S8E, Table S1**). This is accompanied by a significant reduction of the RTC’s forward processivity, defined as the number of nucleotides incorporated by the RTC before the first reversal event (**Figure 2B**). As we increase the concentration of RDV-TP up to 500 µM, we observe a further increase in the reversal population (**Figure S8E, Table S1**), up to the point where no elongating RTC reaches the end of the template (**Figure 2B**). Characterizing the reversals dynamics using a dwell time analysis (**Figure S8FGH, Materials and Methods**), we notice that a reversal trace presents the same characteristics as a trace resulting from the core RTC elongating on a ssRNA template (*52*). Indeed, as opposed to the dwell time distribution of the forward activity, we note only one gamma distribution in the dwell time distribution of the reversals (**Figure S8F**). Furthermore, in absence of RDV-TP, the probability of the slow pathways significantly decreases (**Figure S8DH, Table S2)**, while the gamma distribution characteristic timescale is now similar to the VFNA of the forward trace (**Figure S8CG, Table S2)**. The addition of RDV-TP progressively increases the SNA, VSNA, and LLP probability by ~5-, ~7- and ~11-fold at 500 µM RDV-TP, while the FNA timescale increased slightly (~1.5-fold) without affecting the SNA and VSNA timescales (**Figure S8GH**). This further supports that reversals result from polymerase elongation. In conclusion, RDV-TP incorporation strongly stimulates reversals mediated by nsp13-helicase, decreasing the RTC forward processivity.

### Molnupiravir incorporation also leads to more frequent RTC reversals

We then investigated the mechanism of action of MPV-TP on SARS-CoV-2 RTC using a dsRNA template (**Figure 3**). The addition of 500 µM MPV-TP in the reaction buffer resulted in pauses in the SARS-CoV-2 core RTC elongation traces (**Figure 3A, Figure S9A**). Performing a dwell time analysis, we observed a small (~2-fold) increase in SNA and VSNA timescales, while their respective probabilities remained unaffected (**Figure S9BCD, Table S2**). We also note a ~3-fold increase in the LLP probability (**Figure S9D, Table S2)**. Since MPV-TP is an excellent substrate for the SARS-CoV-2 nsp12-polymerase (*45*), we likely have multiple successive incorporation of MPV-TP when present at 500 µM, increasing the probability of the long-lived pauses.

**Figure 3:**
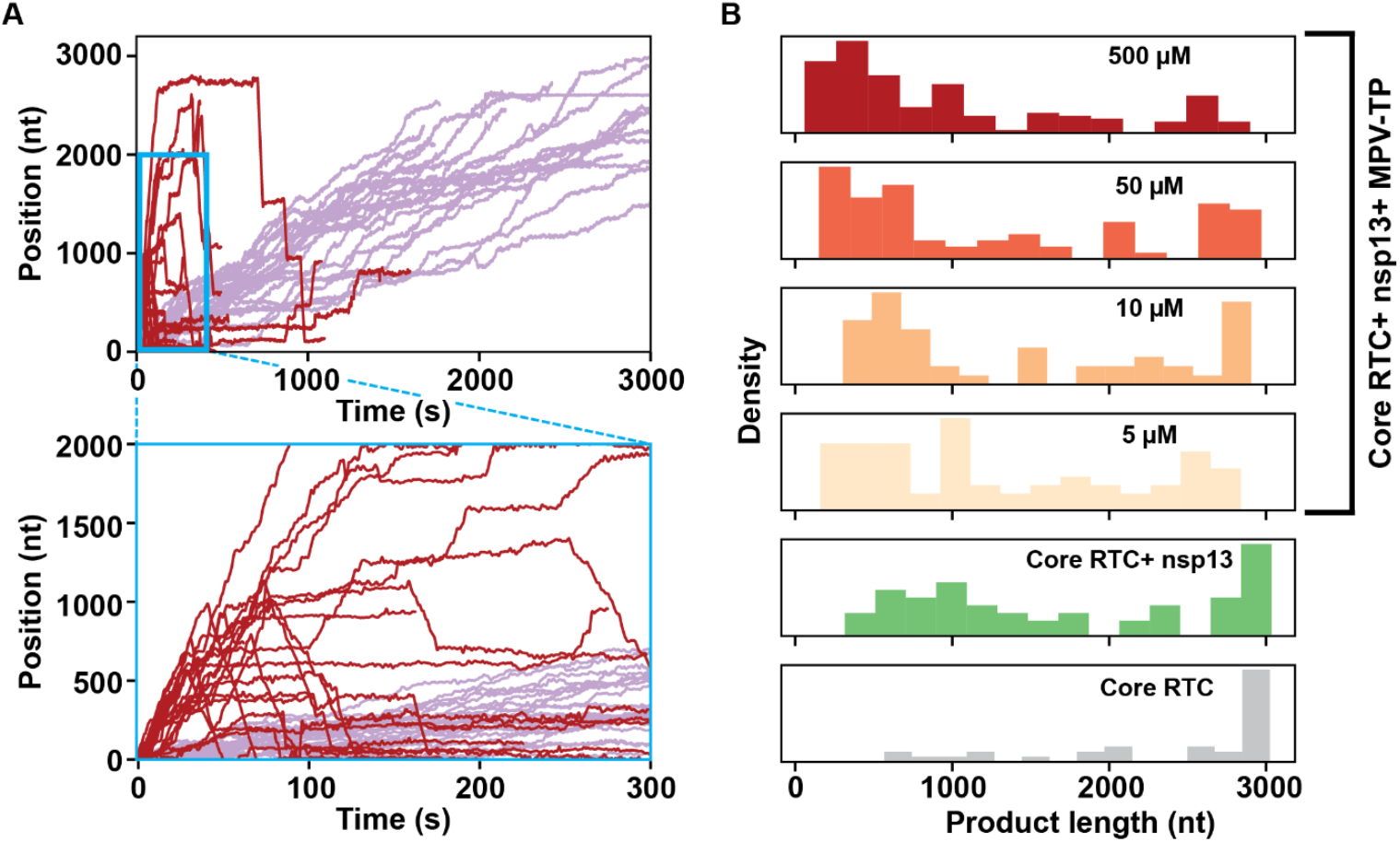
Molnupiravir incorporation increases reversals probability. **(A)** Elongation traces obtained with (dark red) or without (light violet) 20 nM nsp13-helicase, in presence of 500 µM MPV-TP and 500 µM of NTP. The bottom panel shows a zoomed-in view of the elongation traces highlighted with the blue rectangle. **(B)** Histogram of the forward processivity obtained from the elongation traces at different experimental conditions, i.e. without nsp13-helicase (gray), with 20 nM nsp13-helicase (green), with 20 nM nsp13-helicase and the indicated concentration of MPV-TP (5 µM, 10 µM, 50 µM, 500 µM) shown in a gradient from light orange to dark red. All the experiments were performed with 500 µM NTP, at a constant force of 20 pN and 25 °C.

When adding nsp13-helicase together with 500 µM of MPV-TP, the traces largely recovered the elongation velocity measured without MPV-TP (**Figure 3A**). The dwell time analysis showed a mild increase in pause probability in the LLP distribution (**Figure S10ABC, Table S2**). MPV-TP showed no significant difference from CTP when evaluated as a substrate for nsp13-helicase (**Figure S4H**). However, like we observed for RDV-TP, the addition of MPV-TP led to a significant increase of the number of reversals (**Figure 3A**), and their probability, even at low concentrations of MPV-TP (**Figure S10D, Table S1**), i.e. from (0.44 ± 0.13) bp/s to (0.68 ± 0.1) bp/s when increasing MPV-TP concentration from 0 to 10 µM. Raising the concentration further resulted in a decrease of the fraction of RTC reaching the end of the template (**Figure 3B**). The dwell time analysis of the reversals shows an increase in SNA and VSNA probabilities by 4-fold when increasing MPV-TP concentration from 0 to 500 µM, further supporting that the reversals is a product of polymerase elongation (**Figure S10EFG, Table S2**). In conclusion, MPV-TP incorporation significantly increases the probability of the SARS-CoV-2 RTC entering reversals, dramatically reducing the forward processivity of the complex. This result suggests another potential mechanism of action for MPV, in addition to its mutagenic activity.

### Ara-UTP does not induce SARS-CoV-2 RTC reversals

Having re-evaluated the mechanisms of action for RDV and MPV, and their modulation by nsp13-helicase, we aimed to characterize Arabinose (ara)-UTP, for which no mechanism has been reported. Arabinose nucleosides have a different configuration of the 2’ hydroxyl group of their sugar moieties compared to the natural nucleotides and have been used in the treatment of cancer and viral infections (*62, 63*). We investigated how ara-UTP (in competition with UTP) is incorporated by a SARS-CoV-2 core RTC elongating on the ssRNA template (**Materials and Methods, Figure S1B, Figure 4A**) (*33*). A stoichiometry of 1:1 is sufficient to detect ara-UTP incorporation (**Figure S11AB**). Increasing ara-UTP:UTP stoichiometry to 20:1, we observed a significant increase in the number of long-lived pauses interrupting elongation by the core RTC **(Figure 4B)**. Performing a dwell time analysis on the data without ara-UTP and ssRNA template (**Materials and Methods, Figure S11CDE**), we noticed that the LLP are best described by a *t*^−3/2^ power-law distribution (*52*). However, in the presence of ara-UTP, the LLP changes into an exponential (*EXP*_*Ara–UTP*_, **Figure S11CDE, Materials and Methods**), indicating that ara-UTP incorporation induces a single, strong barrier to RTC translocation. From the fit, we note that the FNA, SNA and VSNA timescales remain unchanged, while SNA, VSNA and *EXP*_*Ara–UTP*_ probabilities increased by 3-, 4- and 6.2-fold, respectively, when the stoichiometry of ara-UTP:UTP increased from 1:1 to 20:1 (**Figure S11FG, Table S2**). Repeating the experiments on the dsRNA template (**Figure 1A, Figure S1A, Materials and Methods)** with 10:1 stoichiometry of ara-UTP:UTP, we did not observe any significant impacts of ara-UTP on the elongation velocity (**Figure S12A**). However, we noticed a significant decrease in the mean product length, i.e. (1810± 94) nt vs. (1360 ± 106) nt, indicating that some core RTCs do not recover from the long pauses before the end of the data acquisition (~1h) (**Figure S12B**). In contrast to elongation on ssRNA, the LLP dwell times follow a *t*^−3/2^ power law even in the presence of ara-UTP (**Figure S12C**). From the fit, only the probability of the LLP increases by ~2.6-fold, while all other probabilities and their corresponding timescales remain largely unaffected (**Figure S12DE, Table S2**). In conclusion, ara-UTP incorporation by the core RTC elongating on a dsRNA template induces long-lived pauses.

**Figure 4:**
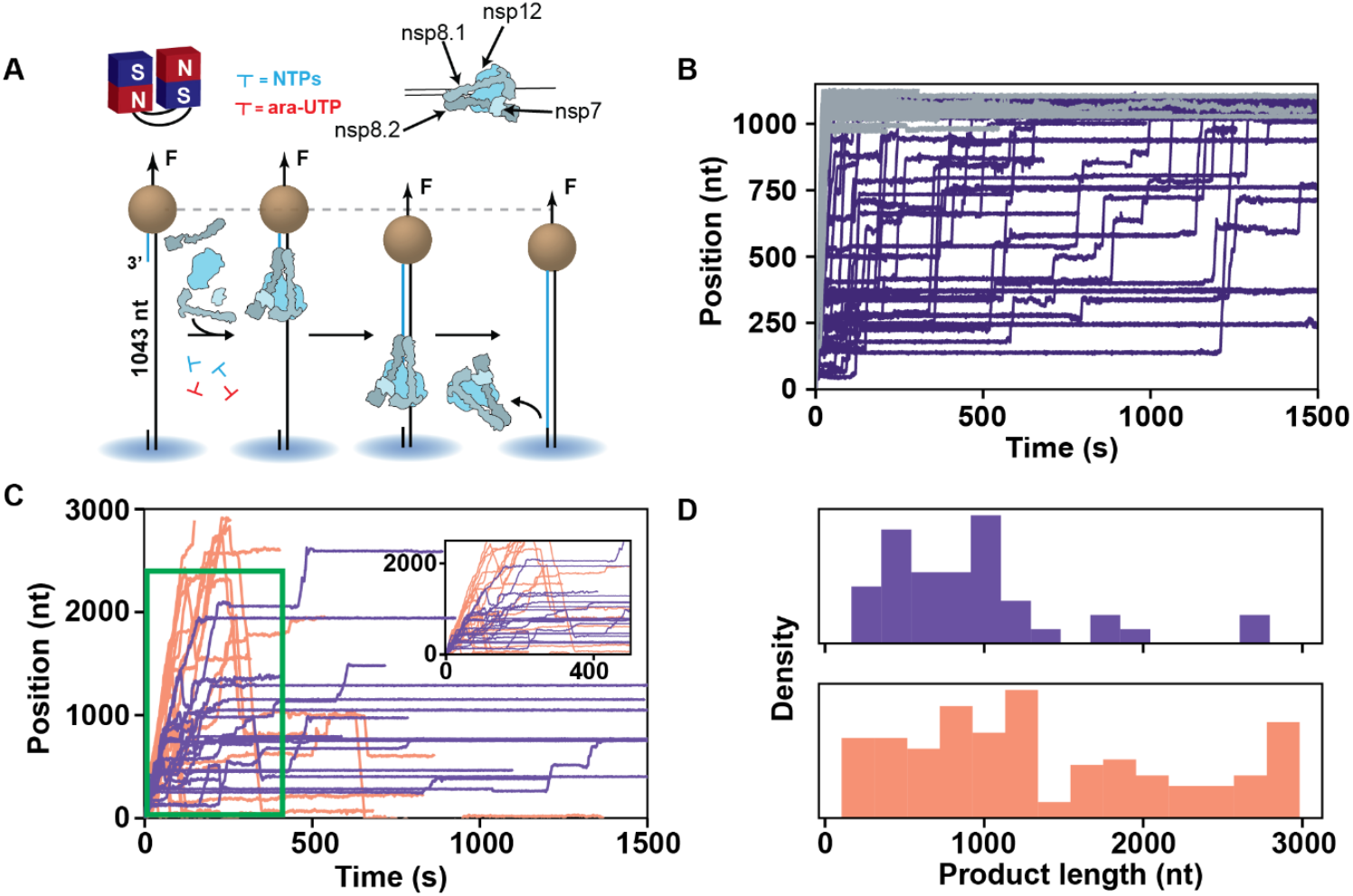
Ara-UTP reduces reversals probability and induces very long pauses during CoV RTC elongation dynamics. **(A)** Schematic of the magnetic tweezers assay to monitor SARS-CoV-2 core RTC RNA synthesis activity on a ssRNA template. (**B)** RNA synthesis activity traces with either no ara-UTP and NTPs at 500 µM (grey), or 1000 µM ara-UTP, 50 µM UTP, 500 µM other NTP (dark violet). **(C)** Elongation traces obtained in the presence (violet) or absence (light orange) of a 10:1 stoichiometry of ara-UTP to UTP (500 µM ara-UTP and 50 µM UTP), with 20 nM nsp13-helicase and 500 µM of all other NTP, on dsRNA (**Figure 1A**). The inset shows zoomed-in elongation traces (green rectangle). **(D)** Histogram of the forward processivity extracted from the elongation traces, as presented in **(C)**, are represented with the same color code.

We next evaluated the impact of ara-UTP on the SARS-CoV-2 RTC with a dsRNA template and in the presence of 20 nM nsp13-helicase (**Figure 1A, Figure S1A, Materials and Methods**). Comparing UTP and ara-UTP as a substrate for nsp13-helicase, we observed no difference (**Figure S4J**). Decreasing UTP concentration to 50 µM increased the reversal probability (**Figure S3, Figure 4C**). When adding ara-UTP at 10-fold excess over UTP, we noticed an increase in the number of LLP interrupting elongation; many of these LLP lasted longer than the duration of the experiment (~1h). Concomitantly, we observed a smaller number of reversal events (**Figure 4C**), suggesting that ara-UTP incorporation reduced their probability (**Figure S13D, Table S1**). From the dwell time distributions, we note a ~2- and ~4.5-fold increase in the VSNA and LLP probabilities, whereas all the other probabilities and their corresponding timescales remain unaffected (**Figure S13ABC, Table S2**). From the forward processivity histogram, we further notice that the elongating RTC hardly reaches the end of the template on the dsRNA construct in the presence of ara-UTP (**Figure 4D**).

Comparison of nucleotide effects on RNA synthesis by RTC reveals significant differences. RDV-TP and MPV-TP incorporation in the nascent strand increases the probability of reversals, thus reducing the RTC processivity (**Figure S8E, Figure S10D, Table S1**), whereas ara-UTP produces very long-lived pauses from which the RTC poorly recovers and inhibits reversals.

## Discussion

In this study, we employed high-throughput magnetic tweezers to investigate the role of nsp13-helicase in complex with the CoV RTC and how the mechanism of action of several nucleotide analogs targeting the nsp12-polymerase is affected by the presence of nsp13-helicase. We show here that nsp13-helicase, in addition to an increase in the RTC average RNA synthesis velocity, also induces reversals in RTC elongation. A duplex RNA fork downstream of the RTC and the ATPase activity of nsp13-helicase are required for such reversals to occur. The reversals probability increases with the stability of the downstream duplex RNA fork and with decreasing NTP concentrations (i.e. below nsp12-polymerase saturation level). We propose that the reversals originate from RTC backtracking, enabling polymerase intramolecular template switching followed by a copy-back RNA synthesis, with the RTC utilizing the former product strand as a template (**Figure 1D**). Nsp13-helicase can hydrolyze any NTP and reaches a significantly larger maximum unwinding velocity with purines than pyrimidines, while having a higher affinity for ATP. We also show that that polymerase intramolecular template switching can be induced by antiviral nucleotide analogs remdesivir and molnupiravir, even at low micromolar range and in the presence of saturating NTP concentrations. However, analogs inducing a strong pause upon incorporation, such as ara-UTP, inhibits reversals.

In our attempt to define nsp13-helicase functions within the CoV RTC, we recently showed that nsp13.2 assists replication through duplex RNA by destabilizing the downstream fork (*24*). Furthermore, our data and kinetic modeling suggest that nsp13.1 and nsp13.2 allosterically control each other’s productive activity (*24*). In other words, both helicases cannot simultaneously translocate on their respective RNA strands. In the present work, we show that nsp13-helicase ATPase activity drives CoV nsp12-polymerase intramolecular template switching and copy-back RNA synthesis in vitro, suggesting a new function for a (+)RNA virus helicase in the context of an elongating RTC. Interestingly, the copy-back RNA synthesis traces are very fast, with very few pauses and are well described by a dynamic similar to the core RTC elongating on a ssRNA template (**Figure S8F-G**) (*52*).

Decreasing the concentration of a single NTP below the saturating concentration for nsp12-polymerase (i.e. below 300 µM (*52*)) is sufficient to increase the probability for the RTC to switch template (**Figure S3**), even though all the other NTPs remain at saturation. We have shown in our previous work that such NTP concentration drives nsp12-polymerase into slower nucleotide addition pathways (*49*), a potential intermediate preceding template switching. Furthermore, a stable dsRNA fork downstream of the RTC enables template switching (**Figure 1AB, Figure S2A-D**). We therefore propose a model where nsp13.2 disengages from the non-template RNA, allowing nsp12-polymerase to enter one of the slower nucleotide addition pathways. From there, nsp12-polymerase can enter the backtrack state. Nsp13.1 then pushes the RTC further backward, such that the 3’-end of the RNA product can snap-back and self-anneal, followed by intramolecular template switching and copy-back RNA synthesis (**Figure 5, Figure 1D**). The CoV RTC has evolved to sense specific signals encoded in the genome, such as the transcription regulatory sequences and their flanking secondary structures during sgRNA synthesis (*64*), to induce polymerase template switching. Future work will investigate whether the mechanism we describe here is conserved for sgRNA synthesis.

**Figure 5:**
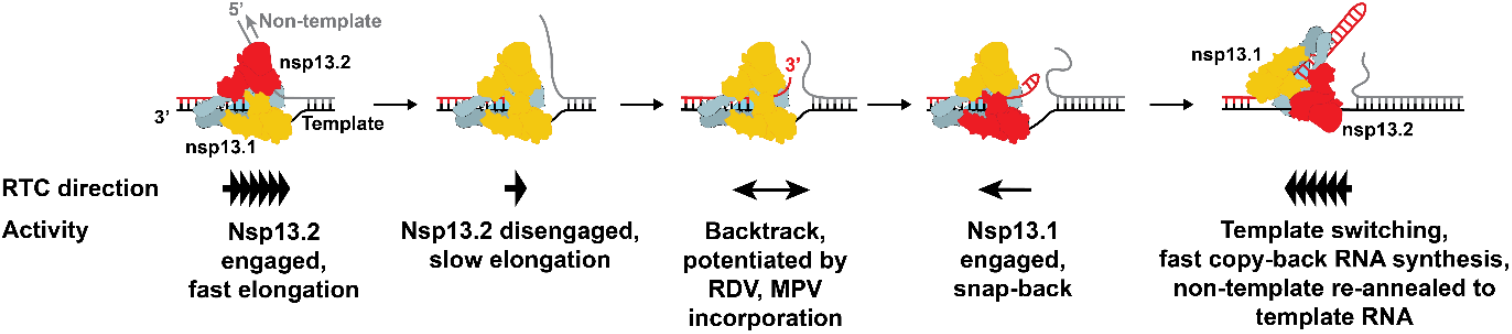
Molecular mechanism of CoV RTC intramolecular template switching and copy-back RNA synthesis. The RTC perform very fast nucleotide addition (VFNA) when nsp13.2 is engaged with the non-template RNA strand. Upon nsp13.2 disengagement, the RTC enters one of the slower nucleotide addition pathways, i.e. SNA or VSNA pathways. From there, the RTC can enter backtrack, which is further potentiated by RDV and MPV, leading nsp13.1 to engage with the template RNA and further push the RTC backward. Thereafter, the product RNA 3’-end snaps back, enabling template-switching and fast copy-back RNA synthesis. This also facilitates the non-template RNA strand to re-anneal to the template RNA. The engagement of either helicase with their respective strand is indicated by the helicase being represented in red.

What could be the function of the RNA produced from copy-back synthesis? It may contribute to genomic diversity through RNA recombination but, given that such RNA would have a hairpin conformation, it is unclear how it may be involved in replication, transcription, or translation. On the other hand, it may increase the pool of defective RNAs (*65*). Such RNA would be particularly inflammatory, as it would be recognized by MDA5 in the infected cell and lead to an innate immune response and interferon production (*65*). Future work will investigate whether such RNA is targeted by nsp15-EndoU to interfere with innate immunity signaling (*66*).

A two-fold mechanism of action has been proposed for RDV. First, RDV-TP induces a polymerase backtracking-related pause upon incorporation (*10, 31–33*). Second, an RDV-MP inserted in the template strand induces a strong pause to the elongating RTC (*47, 49*). The former behavior requires high remdesivir concentration, i.e. above 100 µM, when a physiological NTP concentration is present to result in significant pausing. However, the effective RDV-TP concentration in infected cells is 2-4 µM (*61*) and, at such concentration, the pausing effect is only visible at NTP concentrations three to four orders of magnitude below physiological (0.1 µM) (*10*). In the presence of nsp13-helicase, the post-incorporation pause is only visible in the elongation trace at very high RDV-TP concentration with respect to ATP (500 µM for both, **Figure 2, Figure S8A-D**). Here, we show that having RDV-MP in the template strand does not induce a detectable pause (**Figure S6ABC**). These results argue for an alternative mechanism of action for RDV that differs from inducing a post-incorporation or a template-mediated pause. Indeed, we demonstrate here that RDV-TP incorporation significantly increases reversals probability, even when RDV-TP is in the low micromolar range and in competition with a saturating concentration of ATP, its natural competitor. We previously showed that RDV-TP is incorporated via the slow and very slow nucleotide addition pathways, where it efficiently outcompetes ATP, to induce polymerase backtracking (*49*). Here, we propose the following mechanism to suggest how RDV-TP incorporation induces reversals (**Figure 5**). During RTC elongation, nsp13.2 disengages from the non-template RNA, followed by the RTC entering one of the slower nucleotide addition pathways (SNA, VSNA). There, the RTC incorporates RDV-TP, increasing the probability of polymerase backtracking and the subsequent intramolecular polymerase template switching and copy-back RNA synthesis. Consequently, the incorporation of RDV-TP would decrease the fraction of full-length genomic RNA and concomitantly increase the fraction of defective RNAs. A concentration as low as 10 µM of RDV-TP is sufficient to increase the probability of strand switching from ~40 to ~60% on a 2.9 kbp long genome (**Figure S8E**), while not being detectable in the RTC nucleotide addition kinetics (**Figure S8A**). Extrapolating these numbers to a 29 kbp dsRNA template, i.e. assuming a genome-long dsRNA replication intermediate, the presence of 10 µM RDV-TP would decrease the fraction of RTC reaching the end of such template from 0.6% (in absence of RDV-TP) to 0.01%. This 60-fold decrease in full-length genome production would be a potent mechanism against viral replication.

Furthermore, the strong affinity of nsp13-helicase for ATP (**Figure S5**) and its very high hydrolysis rate even in the absence of RNA (*57*) suggest that nsp13-helicase may deplete ATP in the cell during the CoV infection cycle, further potentiating the polymerase template-switching effect of RDV-TP. Serial passage of SARS-CoV-2 in the presence of RDV resulted in resistance mutations of nsp12-polymerase, i.e. S759A and V792I, with a lower propensity to incorporate RDV-TP than ATP (*67*). Surprisingly, a nsp13-helicase mutant (A336V in SARS-CoV-2) also confers partial RDV resistance to the virus (*68*). We surmise these mutants may have a lower probability of undergoing strand-switching, which poses an interesting avenue for future investigations of resistant mutants.

The mechanism of action of MPV has been proposed to induce mutations post-incorporation in the template strand (*36, 45*). We show here that MPV-TP is very efficiently incorporated by the CoV polymerase and induces pauses upon incorporation (**Figure S9**), which were largely abrogated in the presence of nsp13-helicase (**Figure 3A**). Like with RDV-TP, we also observed a significant increase in reversals probability when increasing MPV-TP concentration (**Figure S10D**). We also attribute these reversals to intramolecular RTC template switching and copy-back RNA synthesis, reducing the forward processivity of the RTC (**Figure 3B**) and increasing the defective RNA production. This would further potentiate MPV therapeutic efficacy, in addition to its demonstrated mutagenic activity (*69*).

While both RDV-TP and MPV-TP stimulates template switching probability, we did not observe such a response with ara-UTP (**Figure 4**). On the contrary, we observed a steep decrease in reversals, and therefore in template switching and copy-back RNA synthesis (**Figure S13D**). Simultaneously, we monitored a strong decrease in the RTC forward processivity when associated with nsp13-helicase, resulting in a premature termination of the elongating RTC. In other words, nsp13-helicase here does not facilitate the RTC escape from the pause induced by ara-UTP incorporation but rather increases the pause’s potency. This suggests that incorporation of ara-UTP by the RTC leads to a stalled complex prior to polymerase template switching, where the RTC remains halted for a long duration, likely because the arabinose moiety prevents further elongation. Our findings indicate that nsp13-helicase may induce unexpected responses from the elongating RTC when incorporating structurally distinct analogs.

Our study further highlights the power of high-throughput magnetic tweezers to shed light on the role of cofactors within an elongating RTC, as well as the mechanisms of action of antivirals. Future research will further expand the CoV RTC with additional factors, such as nsp14-exonuclease (*70*) and investigate whether these factors impact antivirals’ mechanisms of action targeting the CoV RTC. Our bottom-up approach paves the way for a comprehensive understanding of the molecular determinants of coronavirus replication and transcription.

## Acknowledgements

DD was supported by the Interdisciplinary Center for Clinical Research (IZKF) at the University Hospital of the University of Erlangen-Nuremberg, BaSyC – Building a Synthetic Cell” Gravitation grant (024.003.019) of the Netherlands Ministry of Education, Culture and Science (OCW) and the Netherlands Organisation for Scientific Research (NWO), and NWO funding OCENW.XL21.XL21.115. DD, JJA and CEC were supported by grants R01 AI161841-01 and U19 AI171292 from NIAID, NIH. RNK was supported by grant AI158463 from NIAID, NIH. BW and IA were supported by NIGMS R01 GM067153. KDR was supported by NIGMS R35 GM12260. DD thanks Raoul de Groot, Frank van Kuppeveld, Ralph Baric, Mark Denison and Matthias Gotte for fruitful discussions, and Bruno Canard and Etienne Decroly for initial discussions.

## Authors contribution

DD, KDR, CEC and JJA designed the research. DD, AD, AR and SCB designed the experiments. AR, AD and SCB performed the experiments. AD, AR and SCB analyzed the data. PPBA and MK provided support in data acquisition and analysis. JYF, JPB and MV provided remdesivir and oligo-modified remdesivir. FSP and QS made the RNA constructs used for the study. TKA, RNK, BW and IA provided SARS-CoV-2 core RTC proteins. KDR and JCM provided the coronavirus nsp13-helicase. AR, AD and DD interpreted the results and wrote the article. DD supervised the research. All authors have edited the manuscript.

## Supplemental Materials

Materials and Methods Figs. S1 to S13

Table S1 & S2

Supplemental Information can be found online.

