## Supplementary Information for "Template switching by coronavirus polymerase requires helicase activity and is stimulated by remdesivir and molnupiravir"

This document contains Materials and Methods, 13 Supplementary Figures and 2 Supplementary Table.

### Materials and Methods

#### Purification and recombinant protein expression of nsp7, nsp8 and nsp12-polymerase from SARS-CoV-2.

The protocols for expression and purification of nsp7, nsp8 and nsp12-polymerase have been described in detail in Ref. (*1*-*3*).

#### Recombinant Protein Expression of the wild-type and the ATPase dead mutant K288A nsp13-helicases from SARS-CoV-2.

The protocols for expression and purification of the wild-type and the ATPase dead mutant K288A nsp13-helicases have been described in detail in Ref. (*4*).

#### RNA hairpin for polymerase activity fabrication.

The fabrication of the RNA hairpin has been described in detail in Ref. (*5*). The RNA hairpin is made of a 499 bp double stranded RNA stem terminated by a 20 nt loop that is assembled from three ssRNA annealed together (**Figure S1**). The RNA stem is flanked by two spacers, ~800 bp each, containing a biotin- and a digoxigenin-handle, respectively. A gap of 25 nt between the biotin-handle and the hairpin stem serves as the loading site for the polymerase. At forces above 22 pN, the hairpin opens and frees up a 1043 nt ssRNA template for the SARS-CoV-2 polymerase. The RNA construct was synthesized by amplifying DNA fragments in PCR and *in vitro* transcribing them (NEB HiScribe T7 High Yield RNA Synthesis Kit) after purification (Monarch PCR and DNA cleanup kit). ssRNA fragments containing biotin- or digoxigenin-labels were synthesized with biotin-UTP or digoxigenin-UTP (Jena Biosciences) in the reaction. Transcripts were mono-phosphorylated (Antarctic Phosphatase and T4 Polynucleotide Kinase), annealed and ligated. The RNA template sequence is provided in Ref. (*3*).

#### RDV-embedded hairpin for polymerase activity fabrication

The fabrication of the RDV-embedded hairpin is similar to the RNA hairpin fabrication discussed in detail above (**Figure S1**), though the loop oligo sequence includes a remdesivir-monophosphate (MP) such as:
5'-GUAGUGAUUAAGUA**R**ACGAGUAAUCACUACUGGAUCCGUGGGCGCAGCGGAGAAGAA-3',
where **R** is the single Remdesivir-MP(GS-441524-MP). The oligo was synthesized by Dharmacon Inc. and GS-441524 was provided by Gilead Sciences.

#### 2'-O-Me AMP hairpin for polymerase activity fabrication

Similar to the fabrication of the RNA hairpin discussed in detail above (**Figure S1**), with a 2'-O-Me-AMP in the loop oligo sequence such as:
5'-GUAGUGAUUAAGUA**8**ACGAGUAAUCACUACUGGAUCCGUGGGCGCAGCGGAGAAGAA-3',
where **8** is the single 2'-O-Me-AMP residue. The oligo was synthesized by biomers.net GmbH.

**RNA hairpin for helicase activity fabrication**

The helicase RNA hairpin is similar to that of the polymerase RNA hairpin, with the difference that the helicase docking site is on the opposite side of the stem, i.e. on the 5'- end opposite to the 3'- end of the RNA hairpin for polymerase activity, due to the opposite polarity of the helicase and the polymerase. The docking site of the helicase is 20 nt long. In addition, the length of the loop oligo is 4 nt instead of 20 nt for the hairpin for polymerase activity (**Figure S1**).

**dsRNA Construct**

The construct employed here, which has been previously described in detail in Ref. (*5, 6*) and is made of a 4 kb long single-stranded RNA annealed to four ssRNAs: one biotin-labeled strand to attach to the streptavidin-coated magnetic bead, one spacer strand, a ~2.9 kb template strand, and one digoxygenin-labeled strand to attach the tether to the surface glass surface. The template strand ends in 3′ with a small hairpin with the sequence ACGCUUUCGCGT followed by 15 U residues to initiate SARS-CoV-2 RTC or PV RdRp catalyzed RNA synthesis via primer extension.

**Flow cell assembly and surface functionalization.**

The fabrication procedure of the flow cell has been described in detail in Ref. (*7*). To summarize, we sandwiched a double layer of Parafilm by two #1 coverslips, the top one having one hole at each end serving as inlet and outlet, the bottom one being coated with a 0.1% m/V nitrocellulose dissolved in amyl acetate solution. The flow cell is mounted on a custom-built holder and rinsed with ~1 ml of 1x phosphate buffered saline (PBS) solution. 3 μm diameter polystyrene reference beads are non-specifically adsorbed on the bottom coverslip surface by incubating 100 μL of a 1:1000 dilution in PBS (LB30, Sigma Aldrich, stock conc.: 1.828*1011 particles per milliliter) for ~3 min. The tethering of the magnetic beads by the RNA construct relies on a digoxygenin/anti-digoxygenin and biotin-streptavidin attachments at the coverslip surface and the magnetic bead, respectively. Therefore, following a thorough rinsing of the flow cell with PBS, 50 μL of anti-digoxigenin (50 mg/mL in PBS) is incubated for 30 min. The flow cell was rinsed with 1mL of 10 mM Tris, 1 mM EDTA pH 8.0, 750 mM NaCl, 2 mM sodium azide buffer to remove excess of anti-digoxigenin followed by rinsing with another 0.5 ml of 1x TE buffer (10 mM Tris, 1 mM EDTA pH 8.0 supplemented with 150 mM NaCl, 2 mM sodium azide). The surface is then passivated by incubating bovine serum albumin (BSA, New England Biolabs, 10 mg/ml in PBS and 50% glycerol) for 30 min and rinsed with 1x TE buffer.

**Single-molecule RTC elongation activity experiments.**

20 μl of streptavidin-coated Dynal Dynabeads M-270 streptavidin-coated magnetic beads (Thermo Fisher Scientific) was mixed with ~0.1 ng of either RNA hairpin or dsRNA construct (total volume 40 μl) (**Materials and Methods**) and incubated for ~5 min before rinsing with ~2 ml of 1× TE buffer to remove any unbound RNA and the magnetic beads in excess. RNA tethers were sorted for functional hairpins by looking for the characteristic jump in extension of the correct length (~0.6 μm at 30 pN) due to the sudden opening of the hairpin during a force ramp experiment. For the experiments with dsRNA as well, functional tethers were selected with the extension of about (~0.8 – 1 μm at 30

pN). The flow cell was subsequently rinsed with 0.5 ml reaction buffer (50 mM HEPES pH 7.9, 10 mM DTT, 2 μM EDTA, and 5 mM $MgCl_2$). For the experiments with open RNA hairpin, the data acquisition was performed at a force of ~25 pN to keep the hairpin open (*3*), 100 μl of reaction buffer containing 0.6 μM of nsp12, 1.8 μM of nsp7 and nsp8, the indicated concentration of NTPs and of nucleotide analog, i.e. Remdesivir-TP (Gilead Sciences, USA), Molnupiravir-TP (GLPBIO, USA) and Ara-UTP (GLPBIO, USA), were flushed in the flow cell to start the reaction. The experiments were conducted at a constant force for a duration of 30 to 60 minutes. For the experiments with dsRNA, the same concentration of proteins was used, with nsp13-helicase concentration being 20 nM. A constant force of ~20 pN was kept throughout the duration of the experiment (60 minutes). For the experiments without nsp13-helicase, the duration of recording was 90 minutes. The camera frame rate was fixed at 58 Hz the temperature was set to 25°C (*6*). A custom-written LabVIEW routine controlled the data acquisition and the (x-, y-, z-) positions analysis/tracking of both the magnetic and reference beads in real-time (*8*). Mechanical drift correction was performed by subtracting the reference bead position to the magnetic bead position and by applying an autofocus (*3*).

**Single-molecule core RTC + helicase elongation activity experiments on closed hairpin**

The indicated concentrations of M-270 streptavidin-coated magnetic beads and RNA hairpins were mixed and flushed in the flow cell and the subsequent protocol was followed. The only difference is that a force sweep (9, 12, 15, and 20 pN) was done such that the hairpins remained closed. The temperature was set to 25°C. The experiments lasted 60 minutes.

**Single-molecule nsp13-helicase RNA hairpin unwinding experiments**

20 μl of streptavidin-coated Dynal Dynabeads M-270 streptavidin-coated magnetic beads (Thermo Fisher Scientific) was mixed with ~0.1 ng of RNA hairpin for helicase activity. The hairpin selection followed the same protocol as described above (extension of ~0.6 μm at 30 pN). 0.5 ml reaction buffer was flushed followed by 100 μl of reaction mix containing 5 nM nsp13-helicase, desired concentration of NTPs (as stated) at a constant force of 19 pN to keep the hairpins closed. The data was recorded for 60 minutes on a fast sCMOS camera (Dhyana 2100, Tucsen) at 800 Hz image acquisition frequency, subsequently averaged to 400 Hz.

**Data processing**

When using a ssRNA template, i.e. open hairpin, the replication activity of SARS-CoV-2 core RTC converts the tether from ssRNA to dsRNA, which concomitantly decreases the end-to-end extension of the tether. The change in extension measured in micron was subsequently converted into incorporated nucleotides $N_R$ using the following equation (*9*):

$$N_R(F) = N.\frac{L_{ss}(F) - L_{meas}(F)}{L_{ss}(F) - L_{ds}(F)} \quad (1),$$

where $L_{meas}(F)$, $L_{ss}(F)$ and $L_{ds}(F)$ are the measured extension during the experiment, the extension of an ssRNA and of a dsRNA construct, respectively, experiencing a force $F$, and $N$ the number of nucleotides of the ssRNA template.

When using a dsRNA construct, the elongating RTC converts the dsRNA tether into ssRNA, increasing the extension of the tether. To convert the change in extension into a number of incorporated nucleotides $N_R$, we used a modified Equation 1:

$$N_R(F) = N.\frac{L_{meas}(F) - L_{ds}(F)}{L_{ss}(F) - L_{ds}(F)} \quad (2).$$

The traces were subsequently filtered using a Kaiser-Bessel low-pass filter with a cut-off frequency at either 2 Hz or 0.5 Hz for either the open RNA hairpin or the dsRNA and the closed RNA hairpin constructs, respectively. As previously described in Ref. (*9*), a dwell time analysis was performed by scanning the filtered traces with non-overlapping windows of 10 nt to measure the time (coined throughout the manuscript dwell time) for SARS-CoV-2 polymerase to incorporate ten successive nucleotides. The dwell times of all the traces for a given experimental condition were assembled and further analyzed using a maximum likelihood estimation (MLE) fitting routine to extract the parameters from the stochastic-pausing model.

In the case of unwinding experiments with helicase RNA hairpin, the conversion formula used to convert from extension to bp was:

$$N_{bp}(19pN) = \frac{1}{0.000914} * L_{meas}(19\,pN) \quad (3),$$

where $L_{meas}(19\,pN)$ is the measured extension during the experiment. And $N_{bp}(19pN)$ is the number of base pairs unwound. The factor 0.000914 μm/bp was calculated using:

$$\frac{L_{hp-open}(19pN) - L_{hp-closed}(19pN)}{499} = 0.000914 \quad (4),$$

where $L_{hp-open}(19\,pN)$, $L_{hp-closed}(19\,pN)$ are the measured extension of open and closed hairpin respectively at a force of 19 pN. The factor in the denominator is the length of the stem of the hairpin.

For data acquired with SARS-CoV-2 core RTC + ATPase dead mutant K288A nsp13-helicase on a closed helicase RNA hairpin at 9pN force, the data was converted into nucleotides based on the average increase in extension for completed activity traces and low-pass filtered at 1 Hz.

**SARS-CoV-2 RTC forward and reversal trace selection**

From the SARS-CoV-2 RTC nucleotide-converted traces, the forward part of each trace containing reversal was obtained by cutting the trace just before it goes into reversal. And the reversal trace was cut beyond this point.

**SARS-CoV-2 RTC forward processivity and reversal probability analysis.**

The forward processivity of the RTC was determined from the maximum product length preceding the first polymerase reversal event and was then represented into a histogram for each experimental condition.

To calculate the reversal probability, we only considered events where the RTC elongated for a distance longer than 60 nt after the elongation reversed direction. The error bars were estimated from the 95% confidence interval for a binomial distribution.

**SARS-CoV-2 helicase unwinding activity analysis.**

The unwinding velocity of the helicase was determined from the slope of the individual unwinding events, for which only events with a change in extension greater than 30 nm were considered. The unwinding velocities obtained as a function of NTP concentration were fitted with a Michealis-Menten function, extracting the maximum velocity $V_{max}$ and the Michaelis-Menten constant $K_m$ as fitting parameters using the equation:

$$v([NTP]) = \frac{V_{max}[NTP]}{K_m - [NTP]} \qquad (5),$$

where $[NTP]$ is the concentration of the NTP used.

**Maximum likelihood estimation fitting routine.**

The dwell-time distributions were fitted to the experimentally collected dwell-times $\{t_i\}$ by maximizing the log-likelihood function (*10*):

$$LL = \sum_i \ln P_{N\text{nt}}(t_i) \qquad (6),$$

with respect to the characteristic timescales and probabilities. Here $P_{N\text{nt}}$ is the probability of every dwell-time $\{t_i\}$ in the dwell time distribution. We calculated the statistical error on the parameters by applying the MLE fitting procedure on 100 bootstraps of the original data set and reported the standard deviation for each fitting parameter.

**Dwell-time fit-function for nucleotide addition by SARS-CoV-2 core RTC.**

The methodology presented here has been described in detail in Ref. (*11*). The dwell-time distributions of the SARS-CoV-2 core RTC (i.e. in absence of nsp13-helicase) were fitted with a fit-function consisting of one gamma distribution with characteristic timescale $T_{\text{FNA}}$ fitting the peak at short timescale, two exponential distributions with characteristic timescales $T_{\text{SNA}}$ and $T_{\text{VSNA}}$ and a power law distribution of ~$t^{-3/2}$ fitting the long-lived pauses in the dwell-time distributions for longer timescales (*12*):

$$P_{N\text{nt}}(t) \approx \frac{f_{\text{FNA}}}{T_{\text{FNA}}(N-1)!}\left(\frac{tN}{T_{\text{FNA}}}\right)^{N-1} e^{-\frac{tN}{T_{\text{FNA}}}} + Q(t)\left(\frac{f_{\text{SNA}}}{T_{\text{SNA}}} e^{-\frac{t-T_{\text{FNA}}}{T_{\text{SNA}}}} + \frac{f_{\text{VSNA}}}{T_{\text{VSNA}}} e^{-\frac{t-T_{\text{FNA}}}{T_{\text{VSNA}}}} + \frac{f_{\text{LLP}}\sqrt{1+T_{\text{FNA}}}}{2(1+t/1s)^{\frac{3}{2}}}\right) \qquad (7),$$

with $\sum_j f_j = 1$ for $j \in \{\text{FNA}, \text{SNA}, \text{VSNA}, \text{BT}\}$ making the distribution $P_{N\text{nt}}(t)$ is correctly normalized. The regularization function $Q(t) = \frac{(t/T_{\text{FNA}})^{N-1}}{1+(t/T_{\text{FNA}})^{N-1}}$ accounts for the fact that the short timescales are dominated by fast nucleotide addition steps. The cut-off is fixed to the FNA peak position, which is the characteristic timescale of the gamma distribution $T_{\mathrm{FNA}}$. Since the exponential distributions start after the peak of the gamma distribution $T_{\mathrm{FNA}}$, the distributions are also normalized starting from $T_{\mathrm{FNA}}$.

The dwell-time distributions of the SARS-CoV-2 RTC in absence of nsp13-helicase for the experiments with Ara-UTP on ssRNA were fitted with a fit-function consisting of one gamma distribution with characteristic timescale $T_{\mathrm{FNA}}$ fitting the peak at short timescale, three exponential distributions with characteristic timescales $T_{\mathrm{SNA}}$, $T_{\mathrm{VSNA}}$ and , $T_{\mathrm{Ara-UTP}}$ fitting three shoulders in the dwell-time distributions. (*12*)

$$\begin{aligned} P_{N\mathrm{nt}}(t) \approx \frac{f_{\mathrm{FNA}}}{T_{\mathrm{FNA}}(N-1)!}\left(\frac{tN}{T_{\mathrm{FNA}}}\right)^{N-1} e^{-\frac{tN}{T_{\mathrm{FNA}}}} \\ + Q(t)\left(\frac{f_{\mathrm{SNA}}}{T_{\mathrm{SNA}}} e^{-\frac{t-T_{\mathrm{FNA}}}{T_{\mathrm{SNA}}}} + \frac{f_{\mathrm{VSNA}}}{T_{\mathrm{VSNA}}} e^{-\frac{t-T_{\mathrm{FNA}}}{T_{\mathrm{VSNA}}}} + \frac{f_{\mathrm{Ara-UTP}}}{T_{\mathrm{Ara-UTP}}} e^{-\frac{t-T_{\mathrm{Ara-UTP}}}{T_{\mathrm{Ara-UTP}}}}\right) \end{aligned} \quad (8),$$

with $\sum_j f_j = 1$ for $j \in \{\mathrm{FNA, SNA, VSNA, Ara-UTP}\}$ making the distribution $P_{N\mathrm{nt}}(t)$ is correctly normalized. The regularization function $Q(t) = \frac{(t/T_{\mathrm{FNA}})^{N-1}}{1+(t/T_{\mathrm{FNA}})^{N-1}}$ accounts for the fact that the short timescales are dominated by fast nucleotide addition steps. The cut-off is fixed to the FNA peak position, which is the characteristic timescale of the gamma distribution $T_{\mathrm{FNA}}$. Since the exponential distributions start after the peak of the gamma distribution $T_{\mathrm{FNA}}$, the distributions are also normalized starting from $T_{\mathrm{FNA}}$.

For the power law distribution representing a long-lived pause, we have introduced a regularization at 1 s, but the precise timescale does not matter here, as long as it is set within the region dominated by either of the FNA, SNA or VSNA pathways. The power law distribution is cut off with $Q(t)$ and normalized starting from $T_{\mathrm{FNA}}$ like the exponential distributions, following the same rationale. We should, however, be careful in interpreting the long-lived pause probability $f_{\mathrm{LLP}}$, since the power law distribution is only an approximate first passage time distribution and should be interpreted as the relative probability for entering the long-lived pause.

Considering that we observe a clearly separated peak and two shoulders in the dwell-time distributions for the core RTC, we distinguish three characteristic timescales in the dwell-time distribution dominated by fast, slow and very slow nucleotide addition and thus we assume clear separation of timescales for single nucleotide additions $\tau_{\mathrm{FNA}} \ll \tau_{\mathrm{SNA}} \ll \tau_{\mathrm{VSNA}}$. With this assumption, we can derive relations for the characteristic timescales and probabilities in terms of single nucleotide timescales and probabilities as done in (*3*),

$$\begin{gathered} f_{FNA} = p_{\mathrm{FNA}}^N, \quad T_{\mathrm{FNA}} = N\tau_{\mathrm{FNA}}, \\ f_{\mathrm{SNA}} = (p_{\mathrm{FNA}} + p_{\mathrm{SNA}})^N - p_{\mathrm{FNA}}^N, \qquad T_{\mathrm{SNA}} = \frac{Np_{\mathrm{SNA}}(p_{\mathrm{FNA}} + p_{\mathrm{SNA}})^{N-1}}{(p_{\mathrm{FNA}} + p_{\mathrm{SNA}})^N - p_{\mathrm{FNA}}^N}\tau_{\mathrm{SNA}}, \\ f_{\mathrm{VSNA}} = (p_{\mathrm{FNA}} + p_{\mathrm{SNA}} + p_{\mathrm{VSNA}})^N - (p_{\mathrm{FNA}} + p_{\mathrm{SNA}})^N, \\ T_{\mathrm{VSNA}} = \frac{Np_{\mathrm{VSNA}}(p_{\mathrm{FNA}} + p_{\mathrm{SNA}} + p_{\mathrm{VSNA}})^{N-1}}{(p_{\mathrm{FNA}} + p_{\mathrm{SNA}} + p_{\mathrm{VSNA}})^N - (p_{\mathrm{FNA}} + p_{\mathrm{SNA}})^N}\tau_{\mathrm{VSNA}}, \end{gathered} \quad (9),$$

where $p_{\mathrm{FNA}}$, $p_{\mathrm{SNA}}$ and $p_{\mathrm{VSNA}}$ are the probabilities to exit through the FNA, SNA or VSNA pathways for a single nucleotide addition. Furthermore, we added a relative probability to enter the long-lived pause $p_{\mathrm{LLP}}$ such that $p_{\mathrm{FNA}} + p_{\mathrm{SNA}} + p_{\mathrm{VSNA}} + p_{\mathrm{LLP}} = 1$ and $\sum_j f_j = 1$ for $j \in \{\mathrm{FNA; SNA; VSNA; LLP}\}$.

**Dwell-time fit-function for RNA synthesis on dsRNA by the SARS-CoV-2 RTC.**

The dwell-time distributions for forward elongation dynamics by the SARS-CoV-2 RTC (i.e. core RTC in association with nsp13-helicase) were fitted with a dwell-time fit-function consisting of two gamma distributions with characteristic timescales $T_{\mathrm{VFNA}}$ and $T_{\mathrm{FNA}}$ fitting the two peaks at short timescale, two exponential distributions with timescales $T_{\mathrm{SNA}}$ and $T_{\mathrm{VSNA}}$ fitting two shoulders in the dwell-time distributions and a power law distribution ~$t^{-3/2}$ fitting the fat-tail in the dwell-time distributions for longer timescales.

The dwell-time fit-function for the dwell-time distributions of RTC elongation dynamics on a dsRNA template in presence of active nsp13-helicase reads

$$P_{N\mathrm{nt}}(t) \approx \frac{f_{\mathrm{VFNA}}}{T_{\mathrm{VFNA}}(N-1)!} N \left( {}^{tN}/_{T_{\mathrm{VFNA}}} \right)^{N-1} e^{-{}^{tN}/_{T_{\mathrm{VFNA}}}} + \frac{f_{\mathrm{FNA}}}{T_{\mathrm{FNA}}(N-1)!} N \left( {}^{tN}/_{T_{\mathrm{FNA}}} \right)^{N-1} e^{-{}^{tN}/_{T_{\mathrm{FNA}}}} + Q(t) \left( \frac{f_{\mathrm{SNA}}}{T_{\mathrm{SNA}}} e^{-\frac{t-T_{\mathrm{FNA}}}{T_{\mathrm{SNA}}}} + \frac{f_{\mathrm{VSNA}}}{T_{\mathrm{VSNA}}} e^{-\frac{t-T_{\mathrm{FNA}}}{T_{\mathrm{VSNA}}}} + \frac{f_{\mathrm{LLP}}\sqrt{1+T_{\mathrm{FNA}}}}{2(1+t/1s)^{\frac{3}{2}}} \right) \quad (10),$$

with $\sum_j f_j = 1$ for $j \in \{\mathrm{VFNA; FNA; SNA; VSNA; LLP}\}$ making the distribution $P_{N\mathrm{nt}}(t)$ is correctly normalized and the same regularization function $Q(t) = \frac{(t/T_{\mathrm{FNA}})^{N-1}}{1+(t/T_{\mathrm{FNA}})^{N-1}}$ was used. Since the pauses are sub-dominant under both peaks, the cut-off is fixed to the second gamma peak position, corresponding to the FNA pathway $T_{\mathrm{FNA}}$. Like in the dwell-time fit-function for the core RTC, the two exponential distributions and the power law distribution are cut off by $Q(t)$ and normalised accordingly. The power law distribution also has a regularization at 1 s, since the underlying kinetics are not expected to change in presence of active nsp13-helicase. Like for the fit-function for the core RTC dynamics, the long-lived pause probability should be interpreted as a relative probability.

Considering two peaks and two shoulders could be clearly distinguished in the experimental dwell-time distributions, we obtained four separated characteristic timescales dominated by very fast, fast, slow and very slow nucleotide addition pathways. Therefore, we could also assume clear separation of single nucleotide timescales $\tau_{\mathrm{VFNA}} \ll \tau_{\mathrm{FNA}} \ll \tau_{\mathrm{SNA}} \ll \tau_{\mathrm{VSNA}}$.

The dwell-time distributions for reversal elongation dynamics by the SARS-CoV-2 RTC (i.e. core RTC in association with nsp13-helicase) were fitted with a dwell-time fit-function consisting of one gamma distribution with characteristic timescale $T_{\mathrm{FNA}}$ fitting the one peak at short timescale, two exponential distributions with timescales $T_{\mathrm{SNA}}$ and $T_{\mathrm{VSNA}}$ fitting two shoulders in the dwell-time distributions and a power law distribution ~$t^{-3/2}$ fitting the fat-tail in the dwell-time distributions for longer timescales(*12*).

The dwell-time fit-function for the dwell-time distributions of RTC reversal elongation dynamics on a dsRNA template in presence of active nsp13-helicase reads

$$P_{N\mathrm{nt}}(t) \approx \frac{f_{\mathrm{FNA}}}{T_{\mathrm{FNA}}(N-1)!}\left(\frac{tN}{T_{\mathrm{FNA}}}\right)^{N-1} e^{-\frac{tN}{T_{\mathrm{FNA}}}} + Q(t)\left(\frac{f_{\mathrm{SNA}}}{T_{\mathrm{SNA}}} e^{-\frac{t-T_{\mathrm{FNA}}}{T_{\mathrm{SNA}}}} + \frac{f_{\mathrm{VSNA}}}{T_{\mathrm{VSNA}}} e^{-\frac{t-T_{\mathrm{FNA}}}{T_{\mathrm{VSNA}}}} + \frac{f_{\mathrm{LLP}}\sqrt{1+T_{\mathrm{FNA}}}}{2(1+t/1s)^{\frac{3}{2}}}\right) \quad (11),$$

with $\sum_j f_j = 1$ for $j \in \{\mathrm{FNA}, \mathrm{SNA}, \mathrm{VSNA}, \mathrm{BT}\}$ making the distribution $P_{N\mathrm{nt}}(t)$ is correctly normalized. The regularization function $Q(t) = \frac{(t/T_{\mathrm{FNA}})^{N-1}}{1+(t/T_{\mathrm{FNA}})^{N-1}}$ accounts for the fact that the short timescales are dominated by fast nucleotide addition steps. The cut-off is fixed to the FNA peak position, which is the characteristic timescale of the gamma distribution $T_{\mathrm{FNA}}$. Since the exponential distributions start after the peak of the gamma distribution $T_{\mathrm{FNA}}$, the distributions are also normalized starting from $T_{\mathrm{FNA}}$.

### Supplementary Figures

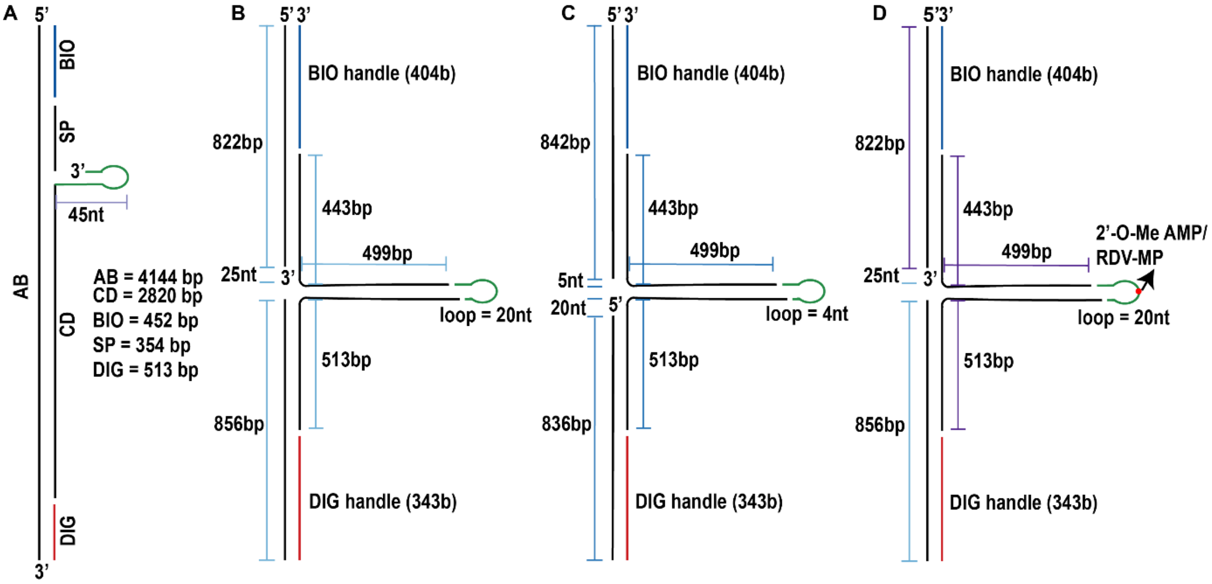

**Figure S1: RNA constructs map. (A)** Schematic representation of the dsRNA construct used in the magnetic tweezers assay. It consists of a 4144 bp non-template strand (AB) annealed to the biotin (BIO) and digoxigenin (DIG) handles, enabling attachment to magnetic beads and the glass surface, respectively. It also contains a spacer (SP) and a 2820 bp template strand (CD) with a 45 nt partialy self complementary 3' overhang (**Materials and Methods**). **(B)** Schematic representation of the RNA hairpin construct used in the magnetic tweezers assay. The construct consists of a 499 bp hairpin flanked by BIO and digoxigenin DIG handles. A 25 nt gap is present between the BIO-handle and the start of the hairpin. (**C**) Schematic representation of the RNA hairpin construct used for nsp13-helicase unwinding activity experiemnts. A 20 nt gap in present at the 5' end between the DIG handle and start of the hairpin. (**D**) Schematic representation of the RNA hairpin construct with a single 2'-O-methylated AMP/RDV-MP insert at the middle of the 20 nt loop of the hairpin indicated by a red dot.

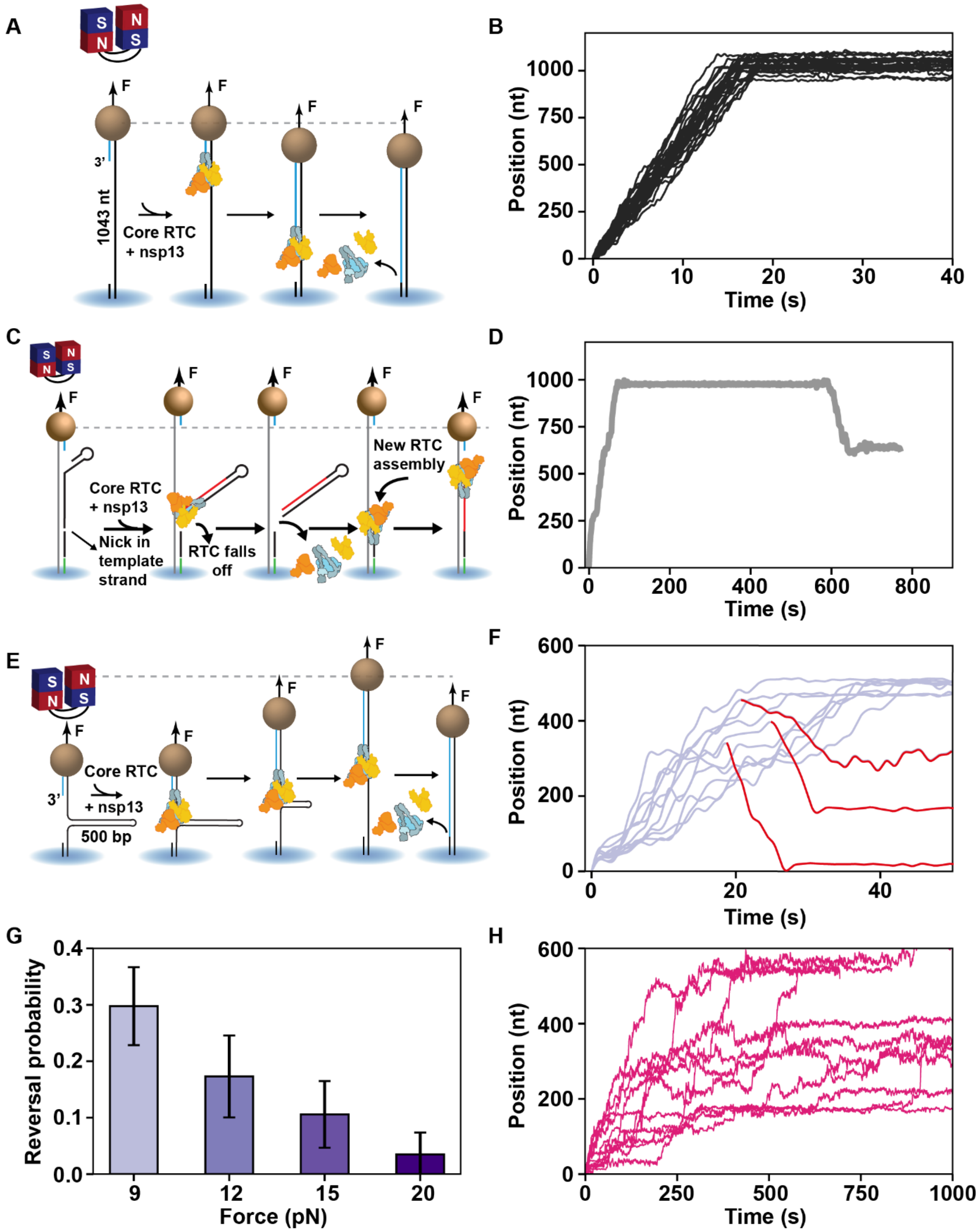

**Figure S2: Nsp13-helicase and duplex RNA are required for reversals to occur. (A)** Schematic of the magnetic tweezers assay to monitor SARS-CoV-2 RTC RNA synthesis activity on a ssRNA template. (**B**) RNA synthesis activity traces in the presence of 500 µM NTPs and 20 nM nsp13-helicase for the experiment described in (A). (**C, D**) Schematic of the magnetic tweezers assay (**C**), and example of an activity trace (**D**), illustrating how a nick in the template strand could be a possible origin of reversals. (**E**) Schematic of the magnetic tweezers assay to monitor SARS-CoV-2 core RTC and nsp13-helicase RNA synthesis on a closed hairpin. (**F**) RNA synthesis activity traces in presence of 1 mM

NTPs and 20 nM nsp13 on closed hairpin at 9 pN force as described in (E). 10 representative activity traces are shown, with the reversals highlighted in red. (**G**) Comparison of the reversal probability as a function of force, extracted from elongation traces acquired with 1 mM NTPs as described in (E). The error bars represent 95% confidence interval. **(H)** Core RTC elongation traces with 20 nM ATPase dead mutant nsp13-helicase (**Materials and Methods**) for the experiment described in (E), 9 pN force and in presence of 1 mM NTPs.

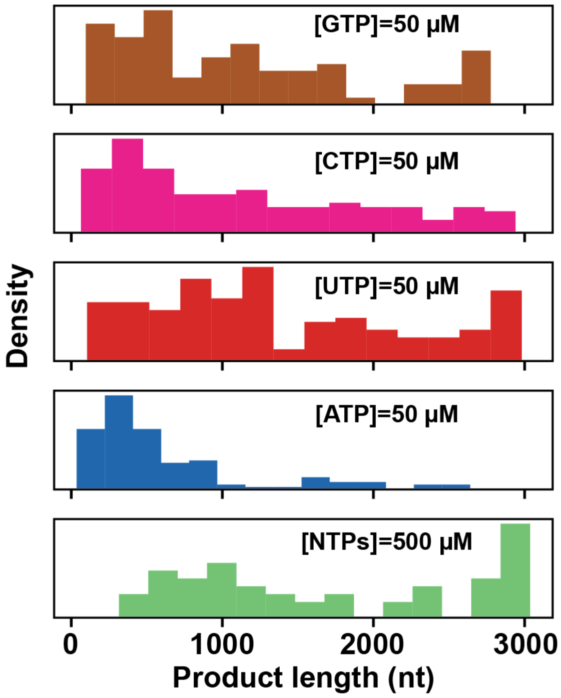

**Figure S3: Forward Processivity of the SARS-CoV-2 core RTC in presence of nsp13-helicase before the first reversal event as a function of NTP concentration.** Histogram of the forward processivity before polymerase template strand switching extracted from the elongation traces of SARS-CoV-2 core RTC and 20 nM nsp13-helicase at different NTP concentrations: 500 μM NTPs (green), 50 μM ATP and 500 μM other NTP (blue), 50 μM UTP and 500 μM other NTP (red), 50 μM CTP and 500 μM other NTP (magenta) and 50 μM GTP and 500 μM other NTP (brown).

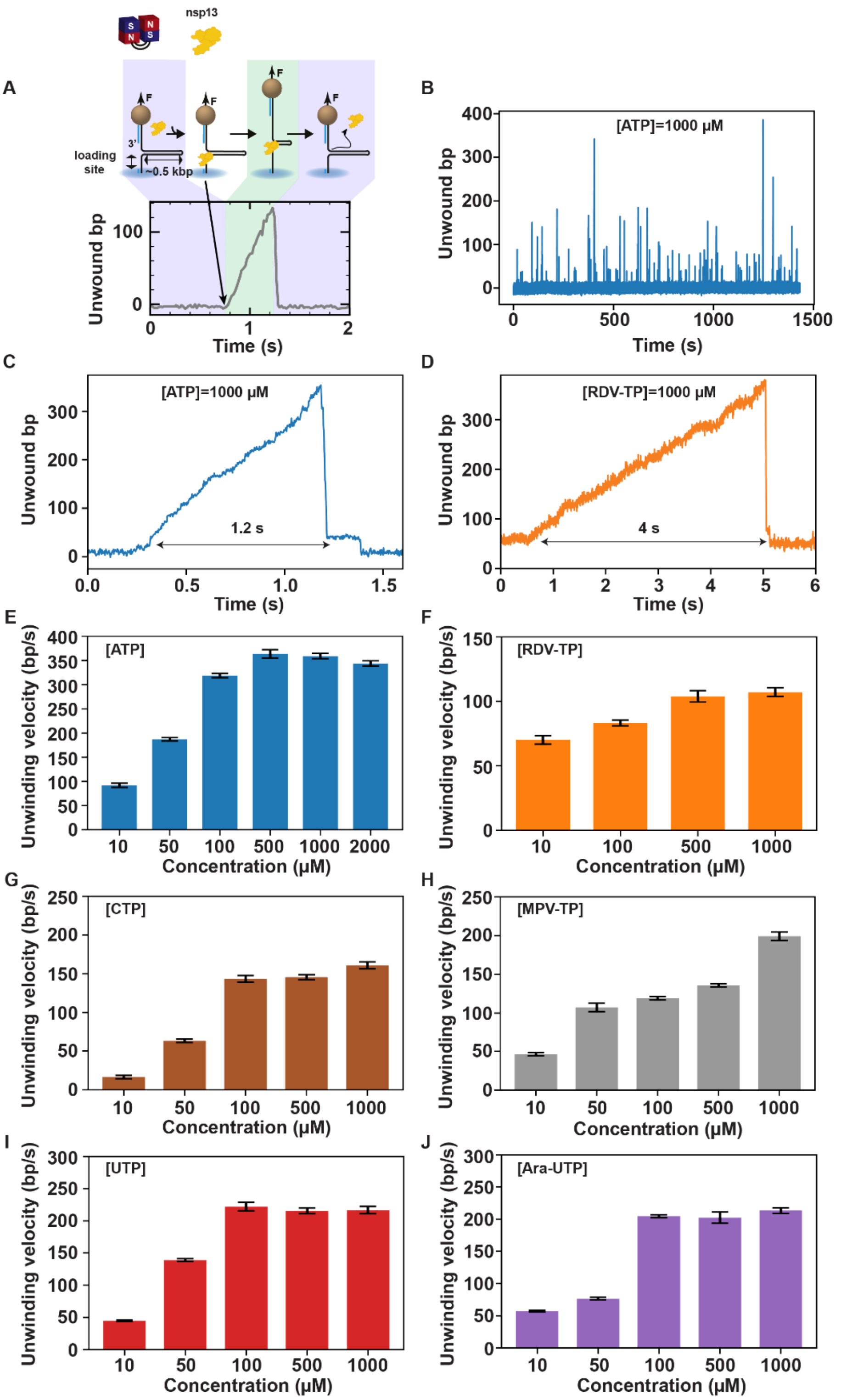

**Figure S4: Comparison of unwinding activity of SARS-CoV-2 nsp13-helicase for different NTPs and nucleotide analogs. (A)** Schematic of the magnetic tweezers assay to monitor SARS-CoV-2 nsp13-helicase activity and an unwinding activity trace below. **(B)** SARS-CoV-2 nsp13-helicase unwinding activity traces, with 1000 μM of ATP and 5 nM nsp13-helicase, where each spike represents an unwinding event. **(C, D)** One single unwinding event obtained with 1 mM of either ATP **(C)** or RDV-TP **(D)**, and 5 nM nsp13-helicase. The time to unwind ~300 bp is also indicated. **(E-J)** Unwinding velocity as a function of nucleotide concentration for different NTPs: ATP **(E)**, CTP **(G)**, UTP **(I)** and their analogs: RDV-TP **(F)**, MPV-TP **(H)**, Ara-UTP **(J)**, extracted from activity traces in the presence of 5 nM nsp13-helicase. The error bars in (**E-J**) are standard error of the mean (SEM). All the nsp13-helicase unwinding activity traces were acquired at 19 pN and 25°C.

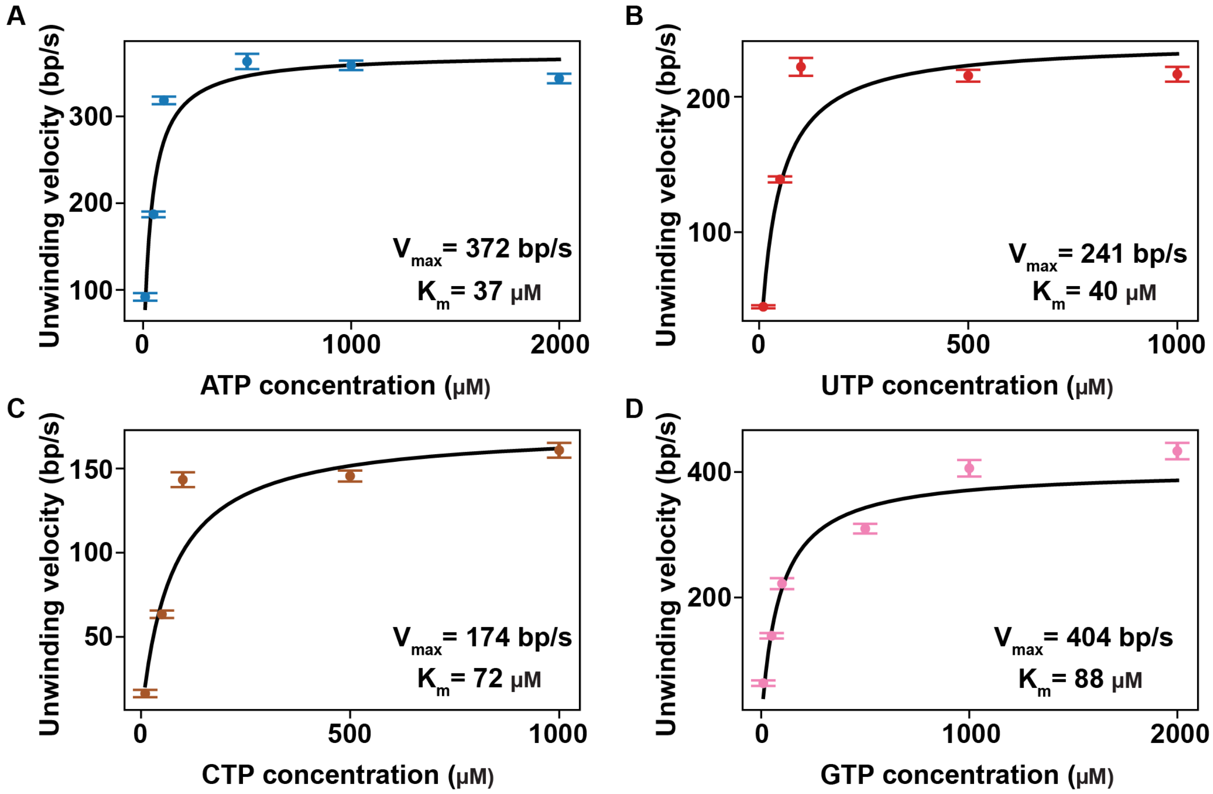

**Figure S5: ATP is the best substrate for SARS-CoV-2 nsp13-helicase. (A-D)** Nsp13-helicase unwinding velocity vs. NTP concentration for either ATP (**A**), UTP (**B**), CTP (**C**) or GTP (**D**). The error bars represent the standard error of the mean. The black solid line is the Michaelis-Menten fit of the unwinding velocity (**Materials and Methods**). The fitting parameters $V_{max}$ and $K_m$ are indicated in each panel.

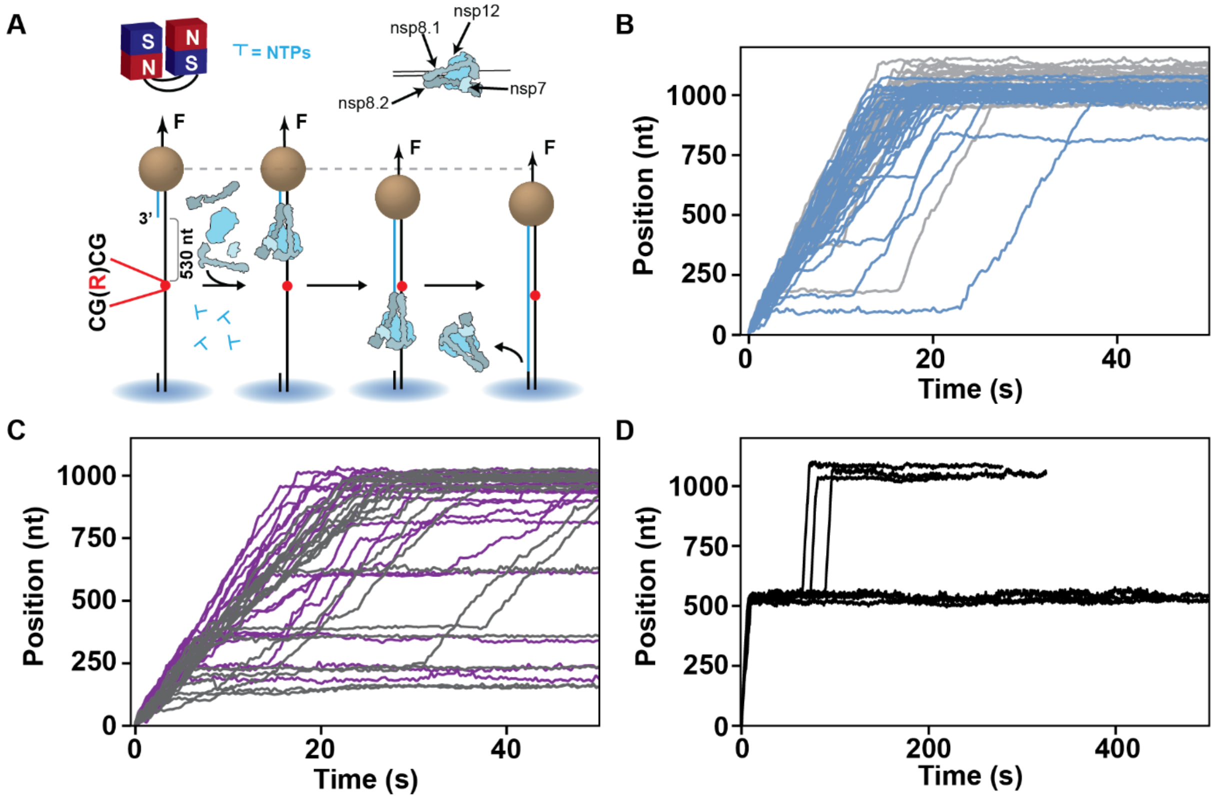

**Figure S6: Template strand inserted RDV-MP does not pause an elongating SARS-CoV-2 core RTC while 2'-O-me AMP does. (A)** Schematic of the magnetic tweezers assay to monitor SARS-CoV-2 core RTC RNA synthesis activity. A magnetic bead is tethered to a glass coverslip by a 1,020-nt long ssRNA template, that includes one single RDV-MP (R) insert 530 nt downstream the 3' end of the primer (blue), which experiences a constant stretching force F of 25 pN. The sequence flanking the RDV-MP insert is indicated. The core RTC assembles at the 3' end of the primer (blue). In the presence of NTP, the core RTC elongates the ssRNA primer, converting the ssRNA template into dsRNA, shortening the tether length. **(B)** SARS-CoV-2 core RTC RNA synthesis activity traces obtained with (blue) RDV-MP inserted ssRNA template or without (light gray), in presence of 500 µM NTPs. **(C)** RNA synthesis activity traces in the presence of 100 µM UTP (violet) or 50 µM UTP (gray) and 500 µM of other NTP, with single RDV-MP insert in the template strand. **(D)** Elongation traces with a 2'-O-methylated AMP insert in the template strand at the same position as RDV-MP in (A) with a concentration of 500 µM NTPs. All RNA synthesis activity traces were acquired at 25 pN and 25°C.

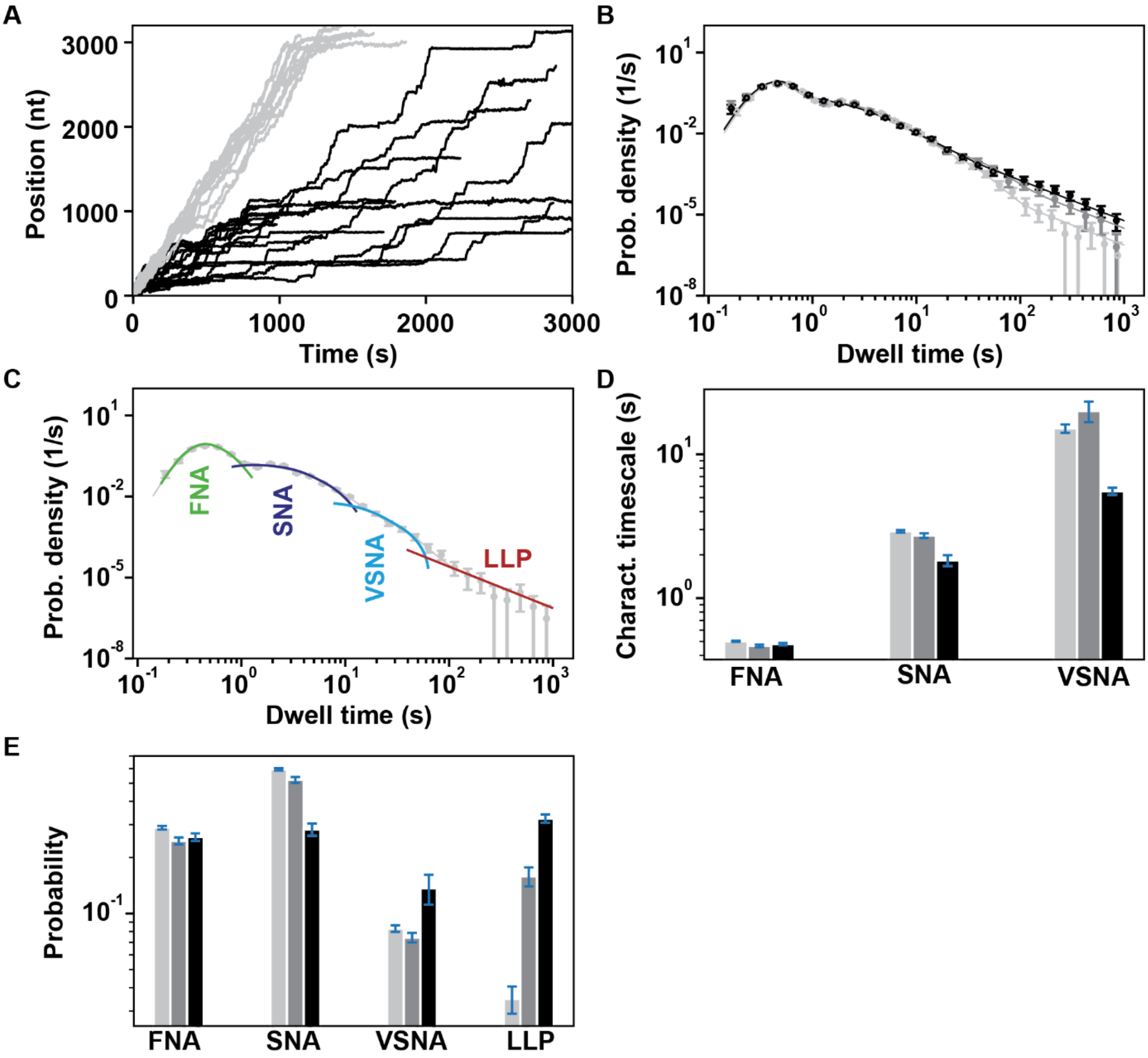

**Figure S7: RDV-TP induces very long-lived pauses upon incorporation by the SARS-CoV-2 core RTC when elongating on a dsRNA template. (A)** RNA synthesis activity traces on dsRNA with (black) or without (light gray) 100 µM RDV-TP, with all other NTPs maintained at a concentration of 500 µM. **(B)** Dwell time distributions of the elongation traces acquired at varying concentration of RDV-TP (0 µM, 20 µM, 100 µM) represented by a color gradient of gray (from light to dark). The solid lines are the corresponding fits (**Materials and Methods**). (**C**)The fit function consists of four probability density functions (pdf's): fast, slow and very slow nucleotide addition (FNA, SNA, VSNA) and long-lived pause (LLP). These pdf's are combined and fitted on the dwell-time distribution (circles) **(D, E)** The fitted parameters, characteristics timescales **(D)** and corresponding probabilities **(E)** for the dwell time distributions presented in **(B)** are represented, with the same reaction conditions and color codes. The error bars represent one standard deviation extracted from 500 bootstrapping. All the experiments were performed at 20 pN and 25°C.

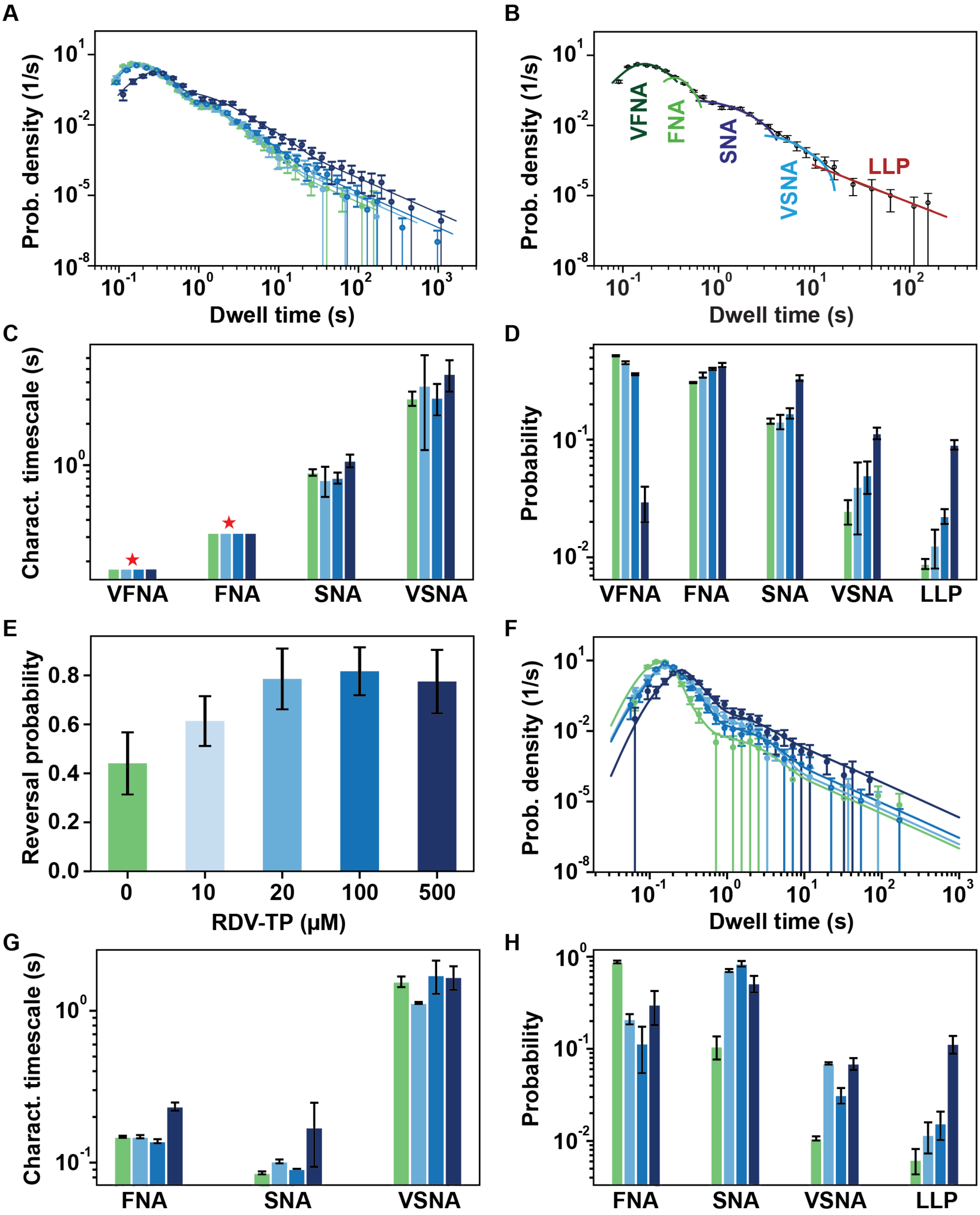

**Figure S8: Comparison of the forward and reversal elongation dynamics of SARS-CoV-2 Core RTC with nsp13-helicase in presence of RDV-TP. (A)** Dwell time distributions (circles) extracted from the RTC forward elongation traces with 20 nM nsp13-helicase at different experimental conditions: without RDV-TP (green) and at varying concentration of RDV-TP (20 µM, 100 µM, 500 µM) shown in a gradient from light blue to dark blue. The solid lines are the corresponding fits (**Materials and Methods**) **(B)** The fit function consists of five probability density functions (pdf's): very fast, fast, slow and very slow nucleotide addition (VFNA, FNA, SNA, VSNA) and long-lived pause (LLP). These pdf's are combined and fitted on the dwell-time distribution (circles) (**Materials and Methods**). **(C, D)** The fitted parameters, characteristics timescales **(C)** and corresponding probabilities **(D)** of the fits to the dwell time distributions presented in **(A)** are represented. VFNA and FNA characteristic time scales in **(C)** were fixed to 0.17 s and 0.32 s respectively and therefore present no error bars (red star). **(E)** Comparison of RTC reversal probability with increasing RDV-TP concentration, extracted from the RTC elongation traces. The errors bars represent 95% confidence interval. **(F)** Dwell time distribution (circles) extracted from the RTC reversal elongation traces at varied reaction conditions, as illustrated in **(A)** and represented with the same color code. The corresponding fits are shown by the solid lines (**Materials and Methods**). **(G, H)** The fitted parameters, characteristics timescales **(G)** and corresponding probabilities **(H)** of the fits to the dwell time distributions presented in **(E)** are represented. The error bars in **(A-D)** and **(F-H)** represent one standard deviation extracted from 100 bootstrapping. All the experiments were performed in the presence of 500 µM NTPs, at 20 pN and 25 °C.

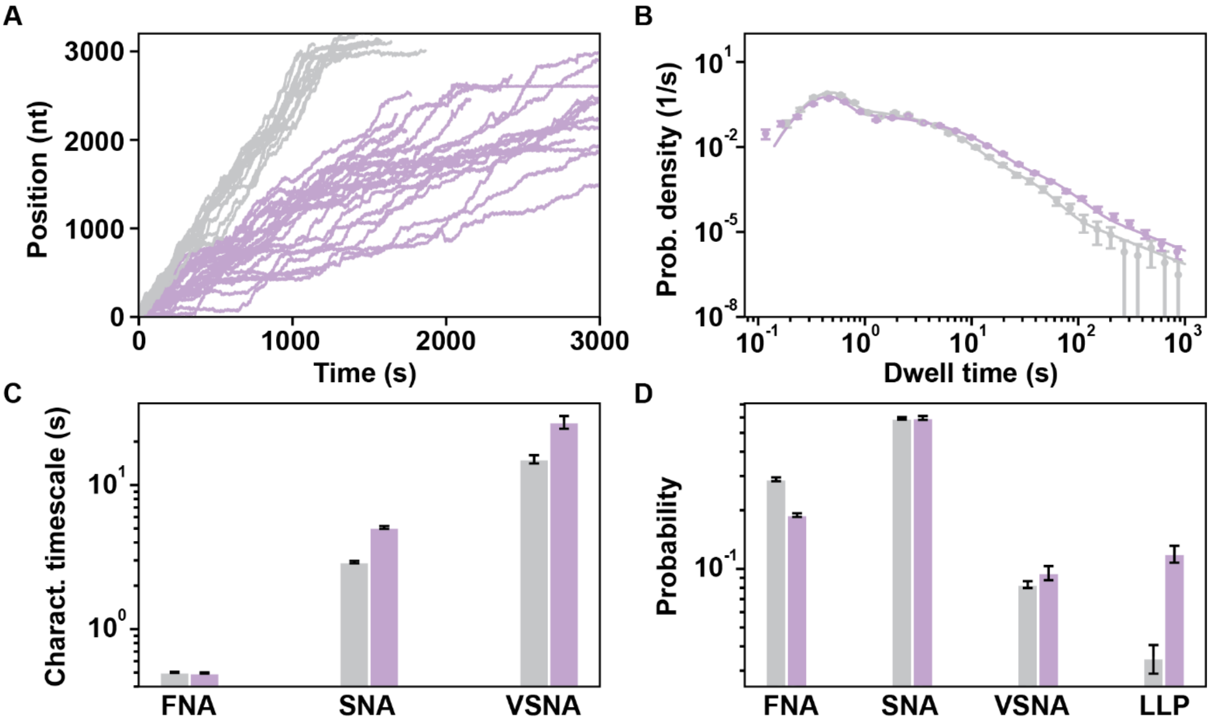

**Figure S9: MPV-TP induces both short and long-lived pauses upon incorporation by the SARS-CoV-2 core RTC on dsRNA. (A)** RNA synthesis activity traces on dsRNA with (light violet) or without (gray) 500 µM MPV-TP, with NTPs maintained at a concentration of 500 µM. **(B)** Dwell time distributions (circles) of SARS-CoV-2 polymerase activity traces under same reaction conditions, as illustrated in **(A)** and represented with the same color codes. The solid lines are the corresponding fits to the dwell time distribution (**Materials and Methods**). **(C, D)** The fitted parameters, characteristics timescales **(C)** and corresponding probabilities **(D)** for the dwell time distributions presented in **(B)** are represented. The error bars are one standard deviation extracted from 500 bootstrapping. All the experiments were performed at 20 pN and 25 °C.

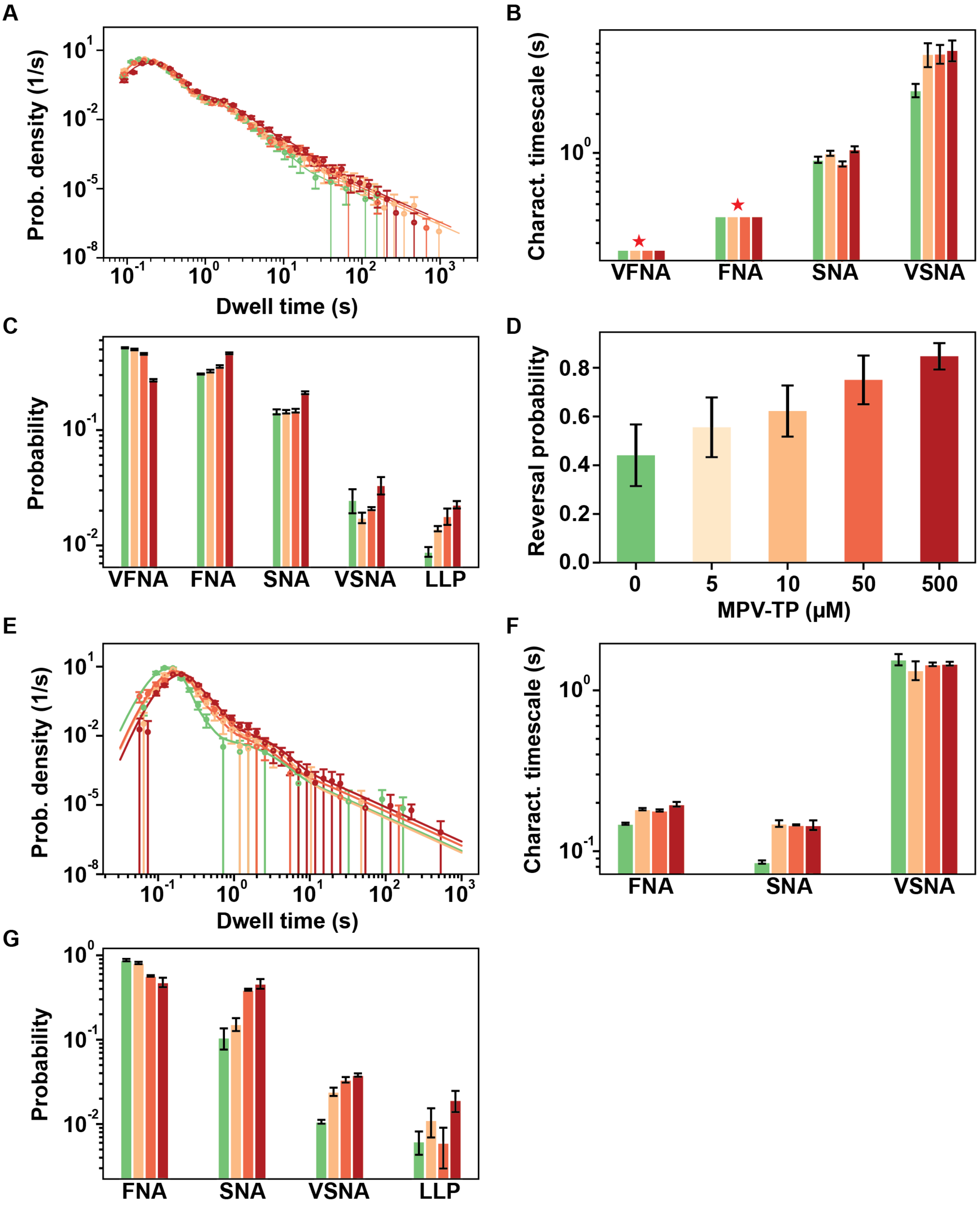

**Figure S10: Comparison of the forward and reversal elongation dynamics of SARS-CoV-2 core RTC with nsp13-helicase in presence of MPV-TP. (A)** Dwell time distributions (circles) extracted from the RTC elongation traces in the forward direction with 20 nM nsp13 at different experimental conditions: without MPV-TP (green) and at varying concentration of MPV-TP (10 µM, 50 µM, 500 µM) shown in a gradient from light orange to dark red. The solid lines are the corresponding fits (**Materials and Methods**). **(B, C)** The fitted parameters, characteristics timescales **(B)** and corresponding probabilities **(C)** of the fits to the dwell time distributions presented in **(A)** are represented. VFNA and FNA characteristic time scales in **(B)** were fixed to 0.17 s and 0.32 s respectively and therefore present no error bars (red star). **(D)** Comparison of RTC reversal probability with increasing MPV-TP concentration, extracted from the RTC elongation traces. The errors bars represent 95% confidence interval. **(E)** Dwell time distribution (circles) extracted from the RTC reversal elongation traces at varied reaction conditions, as illustrated in **(A)** and represented with the same color codes. The corresponding fits are shown by the solid lines (**Materials and Methods**). **(F, G)** The fitted parameters, characteristics timescales **(F)** and corresponding probabilities **(G)** of the fits to the dwell time distributions presented in **(E)** are represented. The error bars in **(A-C)** and **(E-G)** represent one standard deviation extracted from 100 bootstrapping. All the experiments were performed in the presence of 500 µM NTPs, at 20 pN and 25 °C.

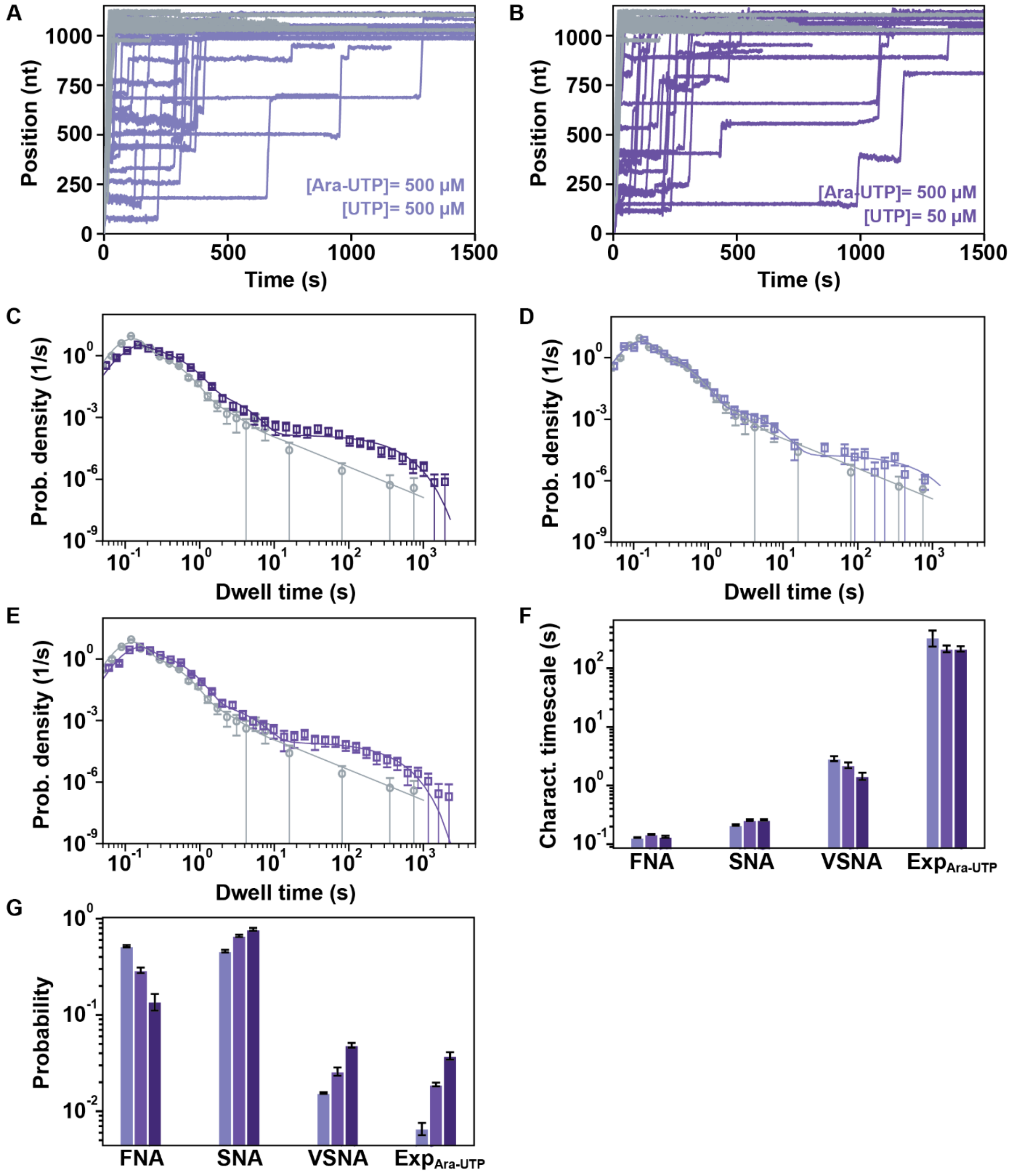

**Figure S11: Effect of ara-UTP on the elongation dynamics of SARS-CoV-2 core RTC on ssRNA.** **(A, B)** RNA synthesis activity traces for **(A)** either 0 μM (gray) or 500 μM (light violet) ara-UTP and 500 μM NTPs, and **(B)** 500 μM ara-UTP, 50 μM UTP and all other NTP at 500 μM (violet). **(C)** Dwell time distributions (circles) extracted from the SARS-CoV-2 core RTC activity traces with either 0 μM ara-UTP and NTPs at 500 μM (grey), or 1000 μM ara-UTP, 50 μM UTP, 500 μM other NTP (dark violet). **(D, E)** Dwell time distributions (circles) extracted from the activity traces under the reaction conditions described in **(A, B)**, respectively, presented with the same color code. The solid lines in **(C, D, E)** are the corresponding fits to the dwell time distributions (**Materials and Methods**). **(F, G)** The fitted parameters, characteristics timescales **(F)** and corresponding probabilities **(G)** for ara-UTP:UTP stoichiometries of 1:1, 10:1 and 20:1 are presented with ascending gradient of violet color. The error bars represent one standard deviation extracted from 100 bootstrapping. All the experiments were performed at 25 pN and 25 °C.

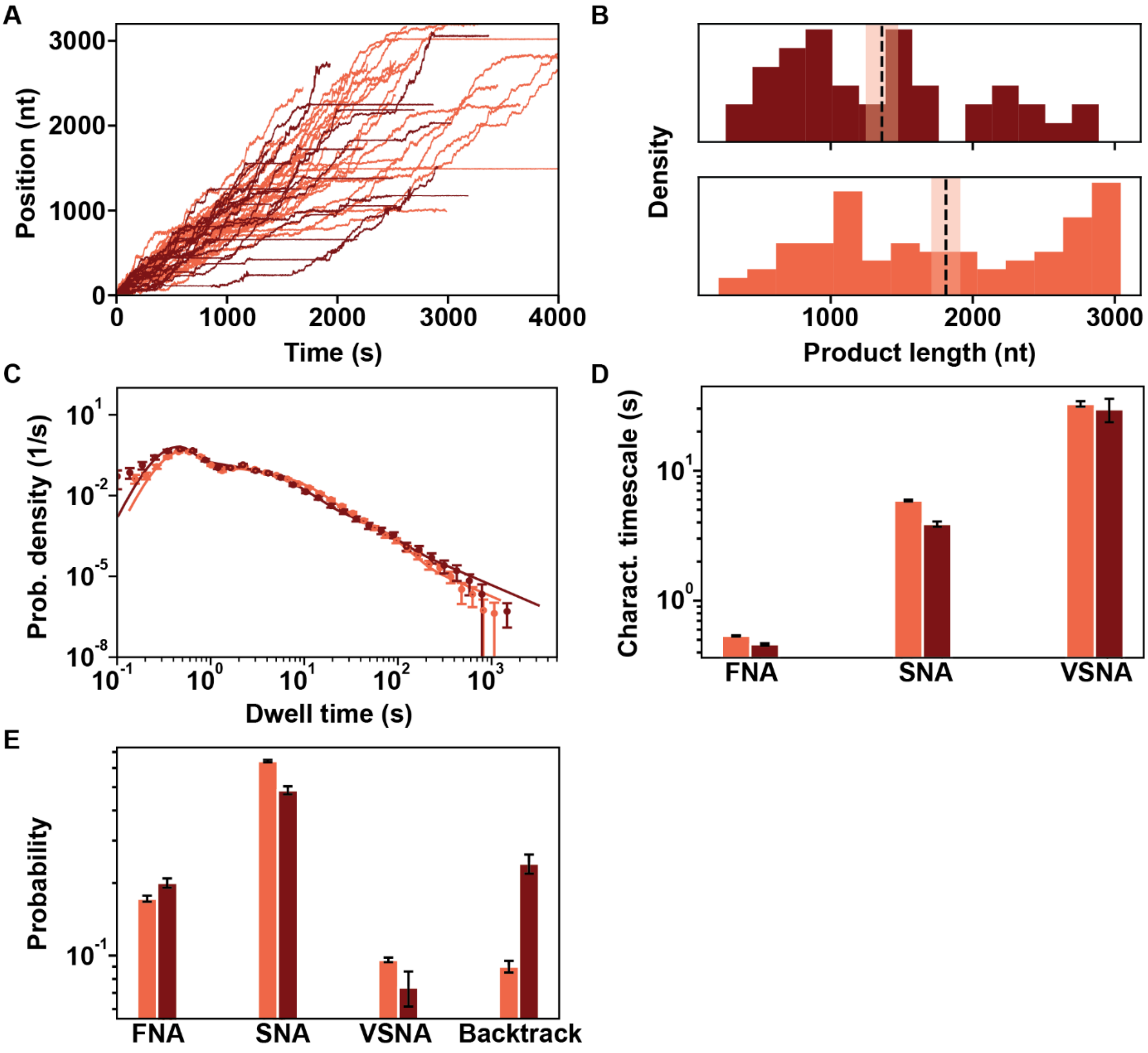

**Figure S12: Effect of ara-UTP on the elongation dynamics of SARS-CoV-2 core RTC on a dsRNA template. (A)** Elongation traces obtained in the presence (dark red) or absence (orange) of a 10:1 stoichiometry of ara-UTP to UTP (500 µM ara-UTP and 50 µM UTP), keeping the other NTP at 500 µM. **(B)** Histogram of the product length extracted from the elongation traces, as presented in **(A)**, are represented with the same color code. The dashed vertical black lines indicate the mean and the shaded area represent one standard deviation extracted from 1000 bootstrapping. **(C)** Dwell time distributions (circles) extracted from the elongation traces, as illustrated in **(A)** are represented with the same color codes. The corresponding solid lines are fits to the dwell time distributions (**Materials and Methods**). **(D, E)** The bar plots represent the characteristics timescales **(D)** and corresponding probabilities **(E)** extracted from the maximum likelihood estimation fits to the dwell time distributions presented in **(C)**. The error bars represent one standard deviation extracted from 100 bootstrapping. All the experiments were performed at 20 pN and 25 °C.

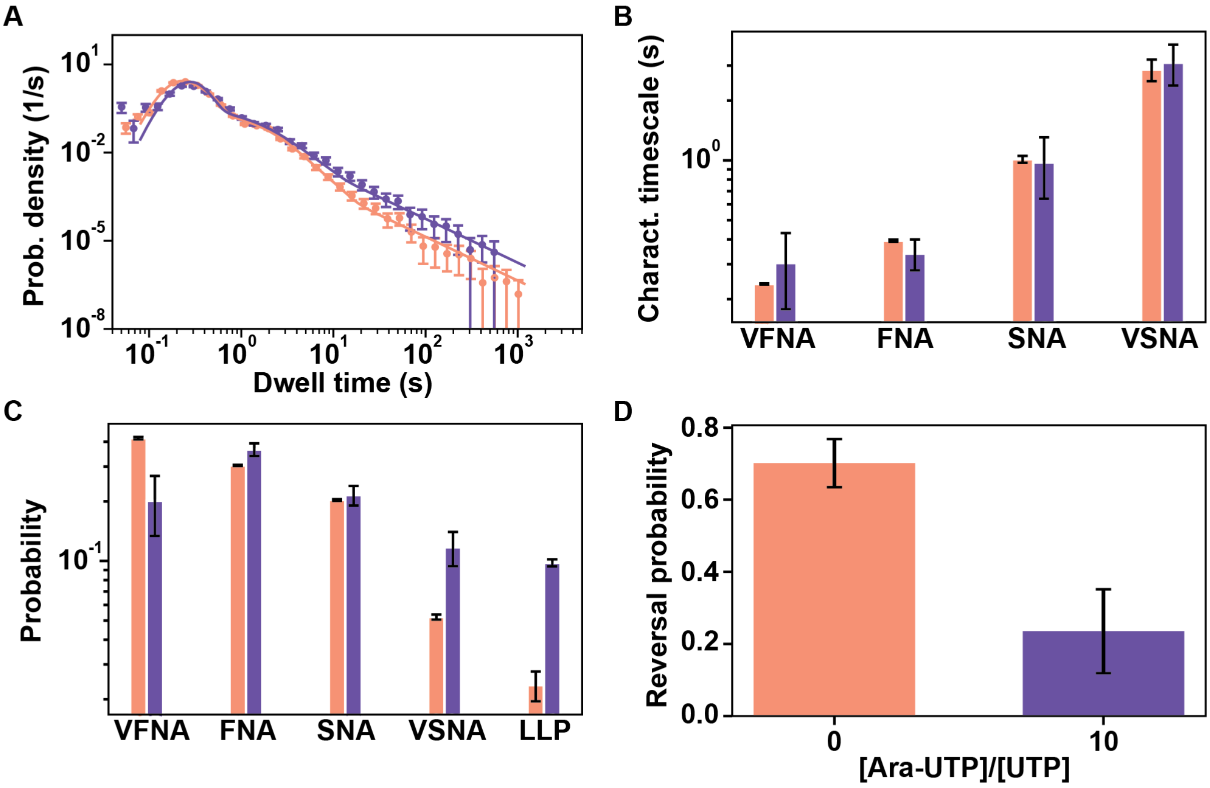

**Figure S13: Effect of ara-UTP on the elongation dynamics of SARS-CoV-2 core RTC and nsp13-helicase on dsRNA. (A)** Dwell time distributions (circles) extracted from the RTC elongation traces with (violet) or without (light orange) 10:1 stoichiometry of Ara-UTP to UTP (500 µM Ara-UTP and 50 µM UTP), 20 nM nsp13-helicase and 500 µM of all other NTP. The corresponding fits are shown by the solid lines (**Materials and Methods**). **(B, C)** The fitted parameters, characteristics timescales **(B)** and corresponding probabilities **(C)** for the dwell time distributions, illustrated in **(A),** are presented with the same color code. The error bars in **(A-C)** represent one standard deviation extracted from 100 bootstrapping. **(D)** RTC reversal probability extracted from the elongation traces, for the same reaction conditions presented in **(A)**, are represented with the same color code. The errors bars represent 95% confidence interval. All the experiments were performed at 20 pN and 25 °C.

**Table S1 :** Summary of the reversal probability for each experimental condition presented in this study

| Construct | nucleotide analog | NTP conc. (µM) | nucleotide analog conc. (µM) | Force (pN) | T (°C) | # traces | # traces with reversals | reversal probability ± error | Figure |
|---|---|---|---|---|---|---|---|---|---|
| ssRNA | NA | 1000 | NA | 9 | 25 | 168 | 50 | 0.29 ± 0.07 | S2E |
| | | | | 12 | | 104 | 18 | 0.17 ± 0.07 | |
| | | | | 15 | | 104 | 11 | 0.1 ± 0.06 | |
| | | | | 20 | | 86 | 3 | 0.03 ± 0.04 | |
| dsRNA | RDV-TP | 500 | 0 | 20 | 25 | 59 | 26 | 0.44 ± 0.13 | S8E |
| | | | 10 | | | 87 | 56 | 0.64 ± 0.1 | |
| | | | 20 | | | 42 | 33 | 0.78 ± 0.12 | |
| | | | 100 | | | 60 | 49 | 0.81 ± 0.09 | |
| | | | 500 | | | 40 | 31 | 0.77 ± 0.13 | |
| dsRNA | MPV-TP | 500 | 0 | 20 | 25 | 59 | 26 | 0.44 ± 0.13 | S10D |
| | | | 5 | | | 63 | 35 | 0.55 ± 0.12 | |
| | | | 10 | | | 82 | 56 | 0.68 ± 0.1 | |
| | | | 50 | | | 72 | 54 | 0.75 ± 0.1 | |
| | | | 500 | | | 170 | 144 | 0.85 ± 0.05 | |
| dsRNA | Ara-UTP | 500 A/C/GTP, 50 UTP | 0 | 20 | 25 | 181 | 127 | 0.70 ± 0.06 | S13D |
| | | 500 A/C/GTP, 50 UTP | 500 | | | 51 | 12 | 0.23 ± 0.11 | |

**Table S2:** Summary of fit parameter statistics for the dwell time distribution for each experimental condition presented in this study

| nucleotide analog | NTP conc. (µM) | nucleotide analog conc. (µM) | T (°C) | Force (pN) | # dwell times | (very fast NA timescale ± std) s | (fast NA timescale ± std) s | (slow NA timescale ± std) s | (very slow NA timescale ± std) s | fast NA probability ± std | slow NA probability ± std | very slow NA probability ± std | long-lived pause probability ± std | Figure |
|---|---|---|---|---|---|---|---|---|---|---|---|---|---|---|
| | | | | | **SARS-CoV-2 core RTC with 20 nM nsp13 RNA synthesis activity traces on dsRNA** | | | | | | | | | |
| | | | | | **forward dwell time distribution** | | | | | | | | | |
| **RDV-TP** | 500 A/C/G/UTP | 0 | 25 | 20 | 11463 | 0.17* | 0.32* | 0.88 ± 0.05 | 3.06 ± 0.36 | 0.31 ± 0.003 | 0.14 ± 0.007 | 0.02 ± 0.006 | 0.009 ± 0.0009 | S8A,C,D |
| | | 20 | | | 6004 | 0.17* | 0.32* | 0.78 ± 0.19 | 3.79 ± 2.50 | 0.35 ± 0.018 | 0.14 ± 0.019 | 0.04 ± 0.024 | 0.013 ± 0.004 | S8A,C,D |
| | | 100 | | | 10943 | 0.17* | 0.32* | 0.81 ± 0.07 | 3.09 ± 0.79 | 0.39 ± 0.009 | 0.17 ± 0.017 | 0.05 ± 0.015 | 0.022 ± 0.003 | S8A,C,D |
| | | 500 | | | 2511 | 0.17* | 0.32* | 1.08 ± 0.11 | 4.61 ± 1.18 | 0.43 ± 0.018 | 0.33 ± 0.018 | 0.11 ± 0.013 | 0.090 ± 0.008 | S8A,C,D |
| **MPV-TP** | 500 A/C/G/UTP | 0 | 25 | 20 | 11463 | 0.17* | 0.32* | 0.88 ± 0.05 | 3.06 ± 0.36 | 0.31 ± 0.003 | 0.14 ± 0.007 | 0.02 ± 0.006 | 0.009 ± 0.0009 | S10A-C |
| | | 10 | | | 12069 | 0.17* | 0.32* | 0.99 ± 0.04 | 5.85 ± 1.23 | 0.32 ± 0.009 | 0.14 ± 0.005 | 0.02 ± 0.002 | 0.014 ± 0.0007 | S10A-C |
| | | 50 | | | 9786 | 0.17* | 0.32* | 0.82 ± 0.04 | 5.89 ± 0.98 | 0.35 ± 0.009 | 0.15 ± 0.005 | 0.02 ± 0.0006 | 0.018 ± 0.0029 | S10A-C |
| | | 500 | | | 15627 | 0.17* | 0.32* | 1.07 ± 0.06 | 6.28 ± 1.15 | 0.46 ± 0.008 | 0.21 ± 0.006 | 0.03 ± 0.006 | 0.022 ± 0.0016 | S10A-C |
| **Ara-UTP** | 500 A/C/GTP, 50 UTP | 0 | 25 | 20 | 21702 | 0.24 ± 0.001 | 0.39 ± 0.005 | 1.01 ± 0.04 | 2.86 ± 0.35 | 0.3 ± 0.002 | 0.2 ± 0.002 | 0.05 ± 0.001 | 0.023 ± 0.004 | S13B-D |
| | | 500 | | | 4415 | 0.3 ± 0.126 | 0.34 ± 0.06 | 0.97 ± 0.33 | 3.09 ± 0.72 | 0.36 ± 0.026 | 0.2 ± 0.024 | 0.12 ± 0.023 | 0.098 ± 0.0039 | S13B-D |
| | | | | | **reversal dwell time distribution** | | | | | | | | | |
| **RDV-TP** | 500 A/C/G/UTP | 0 | 25 | 20 | 1618 | NA | 0.15± 0.002 | 0.09 ± 0.002 | 1.56 ± 0.13 | - | 0.1 ± 0.029 | 0.01 ± 0.0006 | 0.006 ± 0.002 | S8F-H |
| | | 20 | | | 1675 | NA | 0.15 ± 0.004 | 0.1 ± 0.003 | 1.13 ± 0.02 | - | 0.7 ± 0.027 | 0.07± 0.002 | 0.01 ± 0.004 | S8F-H |
| | | 100 | | | 2197 | NA | 0.14 ± 0.004 | 0.09 ± 0.00007 | 1.72 ± 0.42 | - | 0.83 ± 0.059 | 0.03 ± 0.006 | 0.01 ± 0.005 | S8F-H |
| | | 500 | | | 925 | NA | 0.23± 0.015 | 0.17 ± 0.08 | 1.67 ± 0.29 | - | 0.51 ± 0.103 | 0.07 ± 0.001 | 0.11 ± 0.025 | S8F-H |
| **MPV-TP** | 500 A/C/G/UTP | 0 | 25 | 20 | 1618 | NA | 0.15± 0.002 | 0.09 ± 0.002 | 1.56 ± 0.13 | - | 0.1 ± 0.029 | 0.01 ± 0.0006 | 0.006 ± 0.002 | S10E-G |
| | | 10 | | | 920 | NA | 0.18 ± 0.003 | 0.15 ± 0.007 | 1.34 ± 0.18 | - | 0.15 ± 0.027 | 0.02 ± 0.0027 | 0.011 ± 0.004 | S10E-G |
| | | 50 | | | 1764 | NA | 0.18 ± 0.003 | 0.14 ± 0.001 | 1.45 ± 0.04 | - | 0.39 ± 0.011 | 0.03 ± 0.0025 | 0.006 ± 0.003 | S10E-G |
| | | 500 | | | 3677 | NA | 0.19 ± 0.006 | 0.14 ± 0.01 | 1.46 ± 0.04 | - | 0.46 ± 0.060 | 0.04 ± 0.0017 | 0.019 ± 0.005 | S10E-G |
| | | | | | **SARS-CoV-2 core RTC RNA synthesis activity traces on dsRNA** | | | | | | | | | |
| | | | | | **dwell time distribution** | | | | | | | | | |
| **RDV-TP** | 500 A/C/G/UTP | 0 | 25 | 20 | 12973 | NA | 0.5 ± 0.004 | 2.91 ± 0.06 | 15.13 ± 1.0 | - | 0.59 ± 0.009 | 0.08 ± 0.003 | 0.03 ± 0.0006 | S7B-D |
| | | 20 | | | 5501 | NA | 0.46 ± 0.009 | 2.72 ± 0.1 | 19.98 ± 3.2 | - | 0.52 ± 0.018 | 0.07 ± 0.004 | 0.16 ± 0.018 | S7B-D |
| | | 100 | | | 5649 | NA | 0.48 ± 0.009 | 1.83 ± 0.16 | 5.54 ± 0.31 | - | 0.28 ± 0.021 | 0.14 ± 0.025 | 0.32 ± 0.017 | S7B-D |
| **MPV-TP** | 500 A/C/G/UTP | 0 | 25 | 20 | 12973 | NA | 0.5 ± 0.004 | 2.91 ± 0.06 | 15.13 ± 1.0 | - | 0.59 ± 0.009 | 0.08 ± 0.003 | 0.03 ± 0.0006 | S9B-D |
| | | 500 | | | 22608 | NA | 0.49 ± 0.004 | 5.08 ± 0.11 | 27.34 ± 2.7 | - | 0.59 ± 0.013 | 0.09 ± 0.008 | 0.12 ± 0.011 | S9B-D |
| **Ara-UTP** | 500 A/C/GTP, 50 UTP | 0 | 25 | 20 | 16480 | NA | 0.54 ± 0.005 | 5.89 ± 0.09 | 32.63 ± 1.56 | - | 0.64 ± 0.007 | 0.09 ± 0.002 | 0.09 ± 0.005 | S12B-D |
| | | 500 | | | 5637 | NA | 0.46 ± 0.009 | 3.88 ± 0.18 | 29.68 ± 6.03 | - | 0.48 ± 0.018 | 0.07 ± 0.012 | 0.24 ± 0.021 | S12B-D |

| nucleotide analog | NTP conc. (µM) | nucleotide analog conc. (µM) | T (°C) | Force (pN) | # dwell times | (fast NA timescale ± std) s | (slow NA timescale ± std) s | (very slow NA timescale ± std) s | (ara-UTP pause timescale ± std) s | slow NA probability ± std | very slow NA probability ± std | ara-UTP pause probability ± std | long-lived pause probability ± std | Figure |
|---|---|---|---|---|---|---|---|---|---|---|---|---|---|---|
| | | | | | **SARS-CoV-2 core RTC RNA synthesis activity traces on ssRNA** | | | | | | | | | |
| | | | | | **dwell time distribution** | | | | | | | | | |
| **Ara-UTP** | 500 A/C/G/UTP | 0 | 25 | 25 | 14840 | 0.13 ± 0.001 | 0.19 ± 0.008 | 1.09 ± 0.4 | NA | 0.39 ± 0.016 | 0.014 ± 0.002 | NA | 0.007 ± 0.001 | S11C-G |
| | | 500 | | | 7382 | 0.13 ± 0.001 | 0.21 ± 0.007 | 2.88 ± 0.27 | 335.4 ± 102.3 | 0.46 ± 0.014 | 0.015 ± 0.0003 | 0.007 ± 0.0009 | NA | S11C-G |
| | 500 A/C/GTP, 50 UTP | 500 | 25 | 25 | 7313 | 0.15 ± 0.003 | 0.26 ± 0.007 | 2.22 ± 0.25 | 214.8 ± 28.9 | 0.66 ± 0.019 | 0.026 ± 0.002 | 0.019 ± 0.0008 | NA | S11C-G |
| | | 1000 | | | 6320 | 0.13 ± 0.005 | 0.26 ± 0.006 | 1.45 ± 0.21 | 214.5 ± 23 | 0.78 ± 0.027 | 0.048 ± 0.003 | 0.038 ± 0.0033 | NA | S11C-G |

*Very fast and fast NA timescales fixed.
